# The Primacy of Experience in Language Processing: Semantic Priming Is Driven Primarily by Experiential Similarity

**DOI:** 10.1101/2023.03.21.533703

**Authors:** Leonardo Fernandino, Lisa L. Conant

## Abstract

The organization of semantic memory, including memory for word meanings, has long been a central question in cognitive science. Although there is general agreement that word meaning representations must make contact with sensory-motor and affective experiences in a non-arbitrary fashion, the nature of this relationship remains controversial. One prominent view proposes that word meanings are represented directly in terms of their experiential content (i.e., sensory-motor and affective representations). Opponents of this view argue that the representation of word meanings reflects primarily taxonomic structure, that is, their relationships to natural categories. In addition, the recent success of language models based on word co-occurrence (i.e., distributional) information in emulating human linguistic behavior has led to proposals that this kind of information may play an important role in the representation of lexical concepts. We used a semantic priming paradigm designed for representational similarity analysis (RSA) to quantitatively assess how well each of these theories explains the representational similarity pattern for a large set of words. Crucially, we used partial correlation RSA to account for intercorrelations between model predictions, which allowed us to assess, for the first time, the unique effect of each model. Semantic priming was driven primarily by experiential similarity between prime and target, with no evidence of an independent effect of distributional or taxonomic similarity. Furthermore, only the experiential models accounted for unique variance in priming after partialling out explicit similarity ratings. These results support experiential accounts of semantic representation and indicate that, despite their good performance at some linguistic tasks, the distributional models evaluated here do not encode the same kind of information used by the human semantic system.

**Highlights:**

- We used RSA to evaluate three major theories of word meaning representation
- Automatic semantic priming was measured item-wise with high reliability
- Results strongly support representation in terms of experiential information
- Word co-occurrence information did not independently contribute to semantic priming
- RSA and semantic priming can be used to determine the featural content of concepts

**Statement of Relevance:** Understanding the representational code underlying language meaning is not only a central goal of the cognitive sciences but also a gateway to major advances in artificial intelligence and treatment of language disorders. For the first time, we quantitatively assessed the extent to which different kinds of information are encoded in the mental representation of word meanings using an implicit behavioral measure of meaning similarity. We found strong evidence that word meanings encode multimodal experiential information reflecting the functional organization of the brain, in agreement with embodied models of semantics. There was no evidence for distributional information (i.e., derived from patterns of word co-occurrence), indicating that language models such as generative pre-trained transformers (GPTs) do not encode the same kind of information that is represented in human semantic memory. These results indicate that theoretical advancements in this area will require detailed characterizations of how experiential information is implemented in semantic memory.

**Graphical Abstract:** 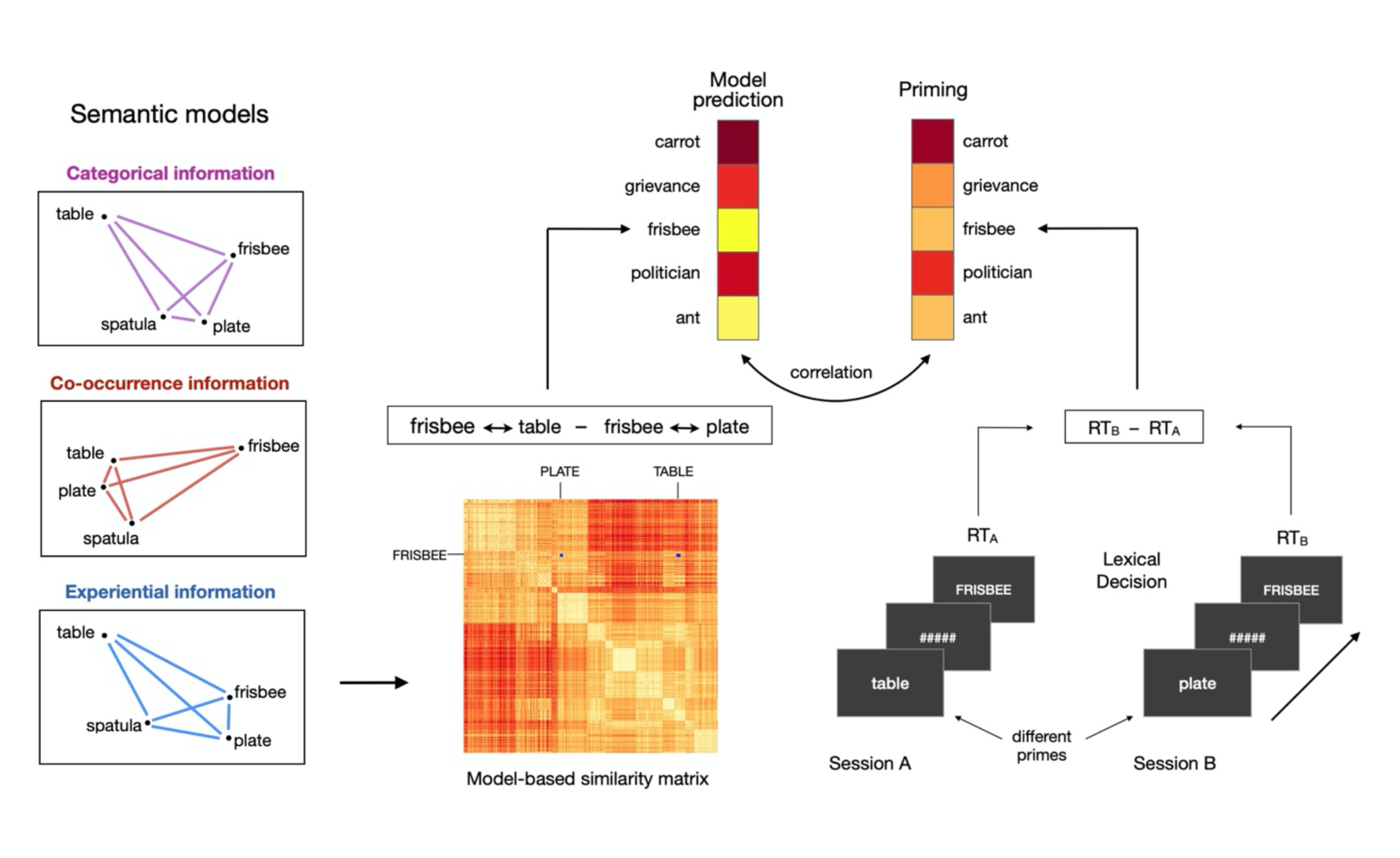

## Introduction

The nature of the relationship between language meaning and the physical reality experienced through the senses has been the subject of scholarly debate since the dawn of the Western intellectual tradition. From the writings of Plato and Aristotle through the works of other luminaries such as Descartes, Locke, Kant, and Wittgenstein, the question of how the meanings of words and sentences are implemented in the mind and brain has figured prominently, and it remains a central issue in the contemporary cognitive sciences (e.g., Barsalou, 1999; Glenberg & Robertson, 2000; Jackendoff, 2002; Rogers & McClelland, 2004; for reviews, see Binder & Fernandino, 2016, and Vigliocco & Vinson, 2007; see also the special issue of *Psychonomic Bulletin and Review*, *23*(4)). Currently, there is consensus among scholars that, aside from physiological processes taking place in sensory and motor organs, any representations or computations underlying language production and comprehension must be somehow implemented in terms of neurophysiological processes in the brain. The ongoing debate concerns the characterization of these language-related neural representations and processes in terms of the kinds of information they encode and the kinds of computations that are performed. Much of this debate has focused on whether, and to what extent, they encode information originating from sensory-motor processes in the course of our interactions with the world (i.e., experiential information), as opposed to information structures that are independent of the organization of the sensory-motor systems.

According to one prominent view, language meaning is represented in the brain as a symbolic system, in which the neural implementation of elementary units of meaning (i.e., lexical concepts) is arbitrarily related to the neural representations and processes involved in perception and action (Fodor, 1980; Newell, 1980; Pylyshyn, 1984; Simon & Newell, 1971). Symbolic-conceptual representations are specified exclusively in terms of other symbolic-conceptual representations, and their correspondence to real-world entities (i.e., their referents) is given via arbitrary associations that remain external to the system. The relationship between conceptual representations and sensory-motor representations is seen as analogous to the relationship between software and hardware in general-purpose computers, in which the former can be completely specified without any reference to the latter. Because a symbol, by definition, is invariant with respect to specific instantiations (i.e., a type does not discriminate between its tokens), symbolic concepts can only represent categories of things, not individual exemplars. Therefore, this view implies that taxonomic structure is the dominant factor in concept representation. In its original formulation this view has been largely discredited because it leads to circular definitions and begs the question of how symbolic representations are ultimately connected to external reality, also known as the symbol-grounding problem (Harnad, 1990; Searle, 1980). Nevertheless, some authors argue that categorical organization still plays a fundamental role in the neural implementation of word meaning (Bedny, 2020; Caramazza & Mahon, 2003; Mahon & Caramazza, 2011).

In contrast to the symbolic view, many authors have argued that experiential information plays an essential role in semantic language processing. They propose that accessing the meaning of a word involves the activation of neural networks storing an ensemble of elementary experiential representations associated with the word form in question (Barsalou, 1999; Damasio, 1989; Glenberg, 1997; Pulvermüller, 1999). These neural networks, and the representations they encode, are thought to overlap with those involved in processing sensory-motor information during perception and action, implying a substantial degree of continuity between sensory-motor and semantic systems. This view, often referred to as “grounded”, “embodied”, or “situated” semantics, is boosted by several lines of evidence pointing to specific interrelationships between language meaning and sensory-motor representations that are not predicted by the symbolic view (for reviews, see Barsalou, Simmons, Barbey, & Wilson, 2003; Binder & Desai, 2011; Hauk & Tschentscher, 2013; Kiefer & Pulvermüller, 2012; Meteyard, Cuadrado, Bahrami, & Vigliocco, 2012).

However, several authors have been skeptical of the idea that experiential information is an essential component of language meaning. It has been argued, for example, that experimental results taken to indicate activation of sensory-motor information during word comprehension may be driven, instead, by the ortho-phonological structure of the corresponding word forms or by their grammatical class (Bedny, Caramazza, Grossman, Pascual-Leone, & Saxe, 2008; Zubicaray, Arciuli, & McMahon, 2013). Another objection is based on the idea that the activation of sensory-motor representations by words and sentences may be epiphenomenal to semantic processing, simply reflecting spreading of activation from conceptual to perceptual and motor representations (Mahon, 2014; Mahon & Caramazza, 2008; Weiskopf, 2010). Some authors cite the existence of category-specific semantic deficits as evidence for a taxonomic, rather than experiential, organization of semantic memory (Caramazza & Mahon, 2003), while others have pointed to the absence of obvious semantic deficits in persons with congenital sensory-motor impairments as a challenge to embodied semantics (Bedny, 2020).

More recently, it has been proposed that statistical patterns of co-occurrence between word forms in natural language may be used by the brain to represent word meaning (Andrews, Frank, & Vigliocco, 2014; Burgess & Lund, 2000; Landauer & Dumais, 1997; Louwerse, 2008; Mandera, Keuleers, & Brysbaert, 2017). This proposal builds on the idea – known as the “distributional hypothesis” – that words with similar meanings tend to occur in similar linguistic contexts, that is, surrounded by similar sets of words. A corollary of this idea is that, given a large enough body of language samples, the degree of overall similarity in meaning between two words can be determined from their respective linguistic contexts (Harris, 1954; Sahlgren, 2008). With the advent of the internet, the availability of extremely large text corpora in digital form have enabled the emergence of high-performance implementations of this idea, known as distributional language models. These models have been shown to approach human-level performance on a number of semantic language tasks (Landauer & Dumais, 1997; Mandera et al., 2017; Pereira et al., 2016), forming the basis of several artificial intelligence applications currently in use.

Distributional structure has been touted as the solution to the problem of lexical semantic representation (Landauer & Dumais, 1997), and it has been argued that the algorithms implemented in distributional semantic models can be considered plausible simulations of how humans learn word meaning (e.g., Landauer & Dumais, 1997; Mandera et al., 2017), although these claims have been disputed (e.g., Glenberg & Robertson, 2000; Perfetti, 1998). It is now generally recognized that a representational system based exclusively on distributional information runs into the same logical pitfall as the original symbolic view, that is, the symbol-grounding problem. Thus, it has been proposed that lexical concepts are represented in terms of a hybrid experiential-distributional code, in which some concepts are directly grounded in experiential information while others are indirectly grounded via their statistical patterns of co-occurrence with experientially grounded concepts (Andrews et al., 2014; Banks, Wingfield, & Connell, 2021; Barsalou, Santos, Simmons, & Wilson, 2008; Louwerse, 2008).

In sum, the current effort to characterize the underlying nature of language meaning can be framed in terms of which information structures must be postulated in order to achieve a neuroscientifically plausible explanation of human semantic behavior and its associated patterns of neural activity. While taxonomic structure is immediately apparent in semantic word processing, there are important reasons (such as the symbol-grounding issue) to believe that lexical concepts are not encoded in terms of symbolic types, and that category-related effects are likely to be an emergent property of the underlying representational code. It is easy to see how taxonomic structure can emerge from the patterns of covariation of experiential features across lexical concepts, since items belonging to the same category tend to be more similar in terms of sensory-motor and affective features than items belonging to different categories (Rosch, Mervis, Gray, Johnson, & Boyes-Braem, 1976; see also Connell, Brand, J., Carney, J., Brysbaert, M., & Lynott, 2019).

Likewise, taxonomic structure could also emerge from distributional structure, since concepts in a given category tend to appear in similar linguistic contexts, as demonstrated by distributional semantic models.

Because experiential similarity structure is also correlated with distributional similarity structure (i.e., concepts with similar experiential features tend to appear in similar linguistic contexts), studies investigating the representational structure of lexical concepts have produced results consistent with both experiential and distributional representations (Louwerse, 2008, 2011). One exception is a recent study that used representational similarity analyses (RSA) of fMRI activation patterns to compare experiential and distributional models in terms of how much unique variance they explained in the similarity structure of two large sets of nouns (Fernandino, Tong, Conant, Humphries, & Binder, 2022). RSA with partial correlations showed that a model based on 48 experiential features (Exp48) accounted for all the variance explained by either of the distributional models tested (word2vec and GloVe) plus an additional 17% of the explainable variance, indicating that experiential information does contribute to the neural activation patterns underlying lexical concepts, with no evidence of specifically distributional information.

Here, we investigated the extent to which lexical concepts reflect experiential, distributional, and taxonomic information using an independent measure of representational similarity, namely, automatic semantic priming in lexical decision (**Figure 1**). Semantic priming has been used extensively to study the organization of semantic memory, and its magnitude has been shown to reflect the degree of semantic relatedness between the prime and the target (Günther, Dudschig, & Kaup, 2016a, 2016b; Heyman, Hutchison, & Storms, 2016; Hutchison, Balota, Cortese, & Watson, 2008; Jones, Kintsch, & Mewhort, 2006). We used RSA to evaluate and contrast 10 models of semantic representation, including two experiential (Exp48 and Lancaster9), five distributional (word2vec, GloVe, fastText, LSA, and HAL) and three taxonomic (WordNet, Taxonomic A, and Taxonomic B) models. Crucially, we took into account the fact that model-based similarity structures are intercorrelated, using partial correlation to evaluate the unique effects of each kind of information. Since, in principle, priming can be driven by semantic similarity (e.g., mouse – rat) as well as by thematic/contextual association (e.g., mouse – cheese) (Hutchison, 2003; Lucas, 2000; Thompson-Schill, Kurtz, & Gabrieli, 1998), we also collected independent ratings of similarity and association for all prime-target pairs. These ratings were used to evaluate (1) the unique contributions of semantic similarity and thematic association (as assessed via explicit ratings) to priming, (2) how well each factor predicts priming relative to the semantic models and (3) how well each model predicts priming when the variance explained by either set of ratings is partialled out.

**Figure 1.**
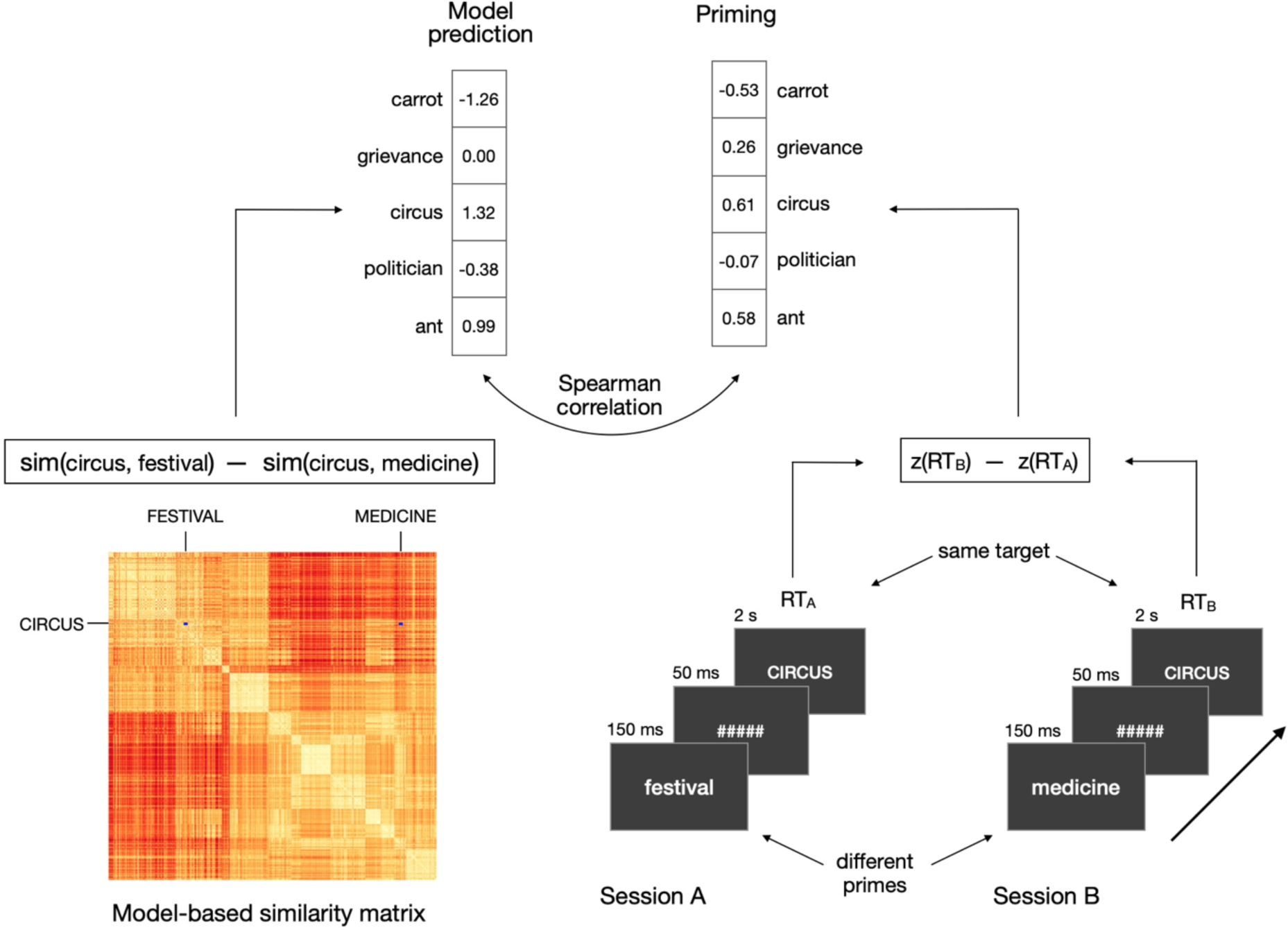
Schematic illustration of the RSA approach. Left: priming predictions were derived from each model of word semantics (only the similarity matrix and predictions from Exp48 are shown). Right: lexical decision was performed on the same set of targets in two testing sessions, yielding a priming value for each target for each participant. Model predictions were evaluated through Spearman correlations with the observed priming.

Finding that a model accounts for semantic priming beyond what can be explained by similarity (or association) ratings indicates that the kinds of information uniquely encoded in the model play a more important role in concept representation than what could be learned from similarity (or association) ratings alone.

While these three kinds of models do reflect distinct representational structures, they also differ in the sense that each kind of model is derived from a different kind of behavior. The experiential models, meant to reflect the sensory-motor-affective organization of the brain, are based on human-provided ratings of the importance of different features of word meaning, which are pre-determined by the investigator. Distributional models, on the other hand, are derived from statistical patterns that emerge from the way people use words to communicate, processed by an algorithm optimized to extract the semantic information implicitly encoded in them. When we evaluate the relative performance of these kinds of models in predicting semantic priming (or any other measure of concept similarity), it is conceivable that differences in performance might be related to differences in information source (i.e., the kind of behavior from which they were derived) rather than the kind of information they encode. At least in principle, it is possible that experiential models, being derived from explicit semantic judgments, have an inherent advantage over distributional models. To investigate this possibility, our analyses included two other models: one based on a set of semantic feature ratings that did *not* reflect the known functional organization of the brain – matching the experiential models in terms of information source but not in terms of information structure – and one based on participant-generated features of lexical concepts, reflecting what people intuitively believe to be the most defining aspects of the meaning of a word (McRae, Cree, Seidenberg, & McNorgan, 2005). The performance of these models can help determine whether representational models based on explicit semantic judgments are inherently superior to distributional models.

## Methods

### Semantic Priming Participants

Thirty-one monolingual English speakers (19 females, ages 22-51, mean age = 31) participated in the study. All were right-handed according to the Edinburgh Handedness Scale, with no history of neurological or psychiatric disorders, and at least 12 years of formal education (mean = 16.5 years). In a simulation-based power analysis, Brysbaert & Stevens (2017) have shown that 6,000 observations (N_obs_ = N_trials_ x N_subjs_) provide enough power (> .8) to detect a 16 ms semantic priming effect in a lexical decision task. With 210 trials (420 paired observations across the two sessions), our study was expected to achieve power above .8 with 29 participants.

### Stimuli

Targets consisted of 210 English nouns and 210 pronounceable pseudowords. Each target was associated with two primes (all real nouns). The task was designed to be performed in two sessions (on different days), such that the same words appeared as targets in both sessions, but each time preceded by a different prime. Targets and primes were selected from the list of 434 nouns for which experiential attribute ratings were available (Binder, Conant, Humphries, Fernandino, Simons, et al., 2016). Based on the results of Hutchison, Balota, Cortese, & Watson (2008), who investigated the impact of several variables on item-level semantic priming, we defined a word’s “priming susceptibility score” as its orthographic neighborhood size (Coltheart’s N; henceforth, OrthN) z-score minus its log-transformed HAL frequency (LogHF) z-score. The 210 nouns with the highest priming susceptibility score were selected as targets (**Supplemental Table 1**). Of these, 157 were concrete and 53 were abstract concepts.

Primes were selected from the 420 nouns with the lowest priming susceptibility score (i.e., relatively high frequency and low OrthN). From this pool of potential primes, we selected two primes for each target based on several criteria. Semantic similarity among words was estimated by averaging together the full pairwise cosine similarity matrix obtained from HAL (http://www.lingexp.uni-tuebingen.de/z2/LSAspaces) and the Wu-Palmer similarity matrix from WordNet (NLTK 3.4.5; https://www.nltk.org). The word with the highest semantic similarity to the target that had not been assigned to a different target was selected as the “close” prime. The “distant” prime was then selected from the list of remaining prime candidates sorted by semantic similarity to the target in descending order. The algorithm searched for the prime candidate with lowest semantic similarity to the target that (a) had not been assigned to a different target, (b) had letter length, number of syllables, LogHF, and OrthN were similar to the close prime, and that (c) was no less then *k* words from the end of the similarity-ranked list. For each new target, *k* was made incrementally larger to select progressively more distant primes, thus providing a wide range of distances from the target. Therefore, rather than selecting primes that were either “related” or “unrelated” to the target (as is common in semantic priming studies), we selected prime pairs whose difference between their similarity to the target (i.e., similarity[target, close prime] – similarity[target, distant prime]) varied continuously across a range (0 to 0.78). None of the primes were strongly associated to their respective targets (all prime-target pairs had a forward association strength lower than 0.1 according to the University of South Florida Free Association Norms).

“Close” and “distant” primes were evenly distributed across two testing sessions, A and B, so that each session contained 105 trials with close primes and 105 trials with distant primes. Cosine similarities between prime and target were matched across the two sessions (session A: mean = 0.400, SD = 0.219; session B: mean = 0.401, SD = 0.206; 94% HDI difference of means: [-.038, .038]; 94% HDI difference of standard deviations: [-.017, .04]). Priming was defined as the difference in standardized response times to the same target between the two sessions (session order was counterbalanced across participants). The primes in the two sessions were matched according to 24 lexical variables, including number of letters, number of phonemes, frequency, orthographic neighborhood size, bigram and trigram frequency, orthographic Levenshtein distance, and age of acquisition (**Supplemental Table 2**).

Pseudowords were created with the MCWord database (https://www.neuro.mcw.edu/mcword) and were matched to the real noun targets in number of letters, bigram and trigram frequency, and orthographic neighborhood size.

### Procedures

Testing was performed over two sessions, approximately one week apart (range: 4-10 days). Each target word was presented once in each session, each time preceded by a different prime (**Figure 1**). The priming effect for a given target word was computed as the difference in z-transformed response times between the two sessions. Each trial started with a central fixation cross (duration jittered 1-2 s), followed by the prime (150 ms), a mask (a sequence of hash marks matching the prime in length, 50 ms), and the target (2 s). The prime was presented in lowercase and the target in uppercase letters. Trials were presented in a different pseudorandomized order for each participant, and the order of presentation of the two primes for a given target (i.e., order of the sessions) was counterbalanced across participants. Two primes (“trust” and “dread”) appeared twice in session 1 and one prime (“excuse”) appeared twice in session 2. All other primes appeared no more than once per session.

Stimuli were presented in light gray with a dark background on the center of a computer screen located 80 cm in front of the participant. Participants were instructed to ignore the prime and make a speeded lexical decision on the target, indicating their response by pressing one of two keys on a response pad with their right index and middle fingers. They were instructed to respond as fast as possible without making mistakes. Stimulus presentation and response recording were performed with E-prime 2.0 software running on a Windows computer and a PST Serial Response Box (Psychology Software Tools, Inc.). For each session, the task was divided into 8 blocks, each lasting 5 minutes. At the beginning of the first session, participants provided informed consent and filled a health history questionnaire. They performed a short practice session (12 trials) immediately before the actual task on both sessions. None of the stimuli included in the experiment appeared in the practice session.

### Similarity and association ratings Participants

Participants were recruited anonymously through the online crowdsourcing service Amazon Mechanical Turk (www.mturk.com) and were financially compensated. They were required to have completed at least 5,000 previous surveys on the Mechanical Turk platform with at least a 95% acceptance rate, and to have account addresses in the United States. All participants reported that they were native speakers of English. Twenty-three participants completed the semantic similarity survey, and 25 participants completed the thematic association survey.

### Stimuli

The stimuli were the same 210 word triplets (i.e., a target and its two respective primes) included in the priming task.

### Procedures

#### Semantic similarity

In each trial, the three words in a triplet were presented simultaneously in a triangular arrangement (**Supplemental Figure 1, left panel**). The word previously used as a target in the lexical decision task appeared at the top center position and its two corresponding primes appeared at the two bottom positions on each side of the screen. A horizontal sliding scale was displayed on the center of the screen. Participants were instructed to position the slider at the point along the scale that best reflected how similar the top word was to the two bottom words. If the top word was equally similar (or equally dissimilar) to the two bottom words, the slider should be positioned in the center of the scale. If the top word was slightly more similar to one bottom word (say, the one on the left) than to the other, the slider should be positioned slightly off center in the direction of the more similar word. The larger the difference in similarity between the two word pairs (i.e., top word-left word similarity versus top word-right word similarity), the further away from the center the slider should be placed. The instructions noted that similarity should be judged based on the extent to which the two concepts had similar properties or could be considered as the same “kind of thing” (e.g., “mouse” and “rat” should be considered highly similar); concepts that often appear together (i.e., thematically or contextually associated, such as “mouse” and “cheese”) but do not have similar properties should not be considered similar.

The task was self-paced, and participants responded by moving the cursor and clicking on a point along the horizontal scale bar on the screen. A circular marker appeared on the scale to indicate the location clicked. Participants could change the position of the marker as many times as desired before clicking on a button labeled “Next” to conclude the trial and start the next one. The response (i.e., the position of the slider) was recorded as a continuous variable, with the center of the scale corresponding to 0, the leftmost position corresponding to -10, the rightmost position corresponding to 10.

#### Thematic association

In each trial, two horizontal scales were displayed on the screen, one on top and the other on the bottom (**Supplemental Figure 1, right panel**). To the left of each scale, two words were presented simultaneously, one being a target in the lexical decision task, the other being one of its two primes. The same target word appeared in both word pairs. For each word pair, participants were instructed to click on a point along the corresponding scale indicating how closely the two words are associated with each other. The instructions noted that association should be judged based on the extent to which the two concepts appear together, not based on whether they have similar properties (e.g., “cake” and “candle” should be considered as strongly associated); concepts that have similar properties but do not often occur in the same context (e.g., “dagger” and “scalpel”) should be considered as weakly or not at all associated. Responses were recorded as a number between 0 (not associated) and 10 (extremely associated). The task was self-paced and both word pairs needed to be rated before participants could proceed to the next trial. Both rating tasks were programmed in Psychopy 3 (Peirce, 2007) and hosted on Pavlovia.org.

#### Data cleaning

Data collected via online crowdsourcing, in which subjects participate anonymously, are more likely to include non-compliant participants. To identify poor quality data (i.e., data from participants who did not follow the task instructions), we computed the Pearson correlation between the ratings from each participant and the group median ratings.

### Semantic models

#### Experiential models

The *Exp48* model is based on the relative importance of 48 dimensions of phenomenological experience chosen to reflect the known functional architecture of the brain. It is based on the experiential feature norms obtained via online crowdsourcing by Binder et al. (2016). Each model dimension encodes the relative importance of an experiential feature to word meaning according to ratings on a Likert-type scale. Exp48 includes perceptual, motor, spatial, temporal, causal, valuative, and goal-related dimensions that can be mapped onto independently established neurocognitive processes (i.e., processes operationalized independently of semantic tasks; see **Supplemental Table 3** for details). The feature ratings stand, roughly, for the extent to which a given functional brain system (e.g., the visual motion system) is activated when a concept is retrieved.

The *Lancaster9* model is based on the Lancaster Sensorimotor Norms (Lynott, Connell, Brysbaert, Brand, & Carney, 2019). Similarly to Exp48, it quantifies the importance of each feature to the meaning of a word based on ratings collected via online crowdsourcing. The norms consist of 11 dimensions corresponding to major sensory and motor modalities of experience: vision, hearing, touch, taste, smell, interoception, hand action, mouth action, foot action, head action, and torso action. Thus, the Lancaster norms contain 5 dimensions of action but only 1 dimension for each of the other sensory modalities. Because semantic similarities based on all 11 dimensions would be disproportionally weighted towards the motor modality, we excluded the Torso Action and Head Action dimensions. The Torso Action dimension seems to be the least important of the five, with the lowest standard deviation and lowest uniqueness score (Lynott et al., 2019, Table 2). Examination of the Head Action dimension in the Lancaster database shows that it conflates at least two very different modalities of experience. Some of the concepts that score highest on this dimension involve motor actions of the head (e.g., “squinting”, “wink”, “blink”, “look back”), while others are instead related to mental and perceptual processes, such as “brainpower”, “thinking”, “memorization”, “vision”, “hearing”, “sight”, and so on. In addition, some of the high-scoring concepts in this dimension are generally related to the head but do not involve any actions (e.g., “nostril”, “hair line”, “eyepatch”, “turban”, “TV”, “headache”). Therefore, these two dimensions were not included in our Lancaster9 semantic model, which consisted of 9 feature ratings.

#### Taxonomic models

*WordNet* (Miller, Beckwith, Fellbaum, Gross, & Miller, 1990) is the largest and most influential database of taxonomic information for lexical concepts. It is organized as a knowledge graph in which words are grouped into sets of synonyms (“synsets”), each expressing a distinct concept. Synsets are interconnected according to conceptual-semantic relations. Our WordNet model is based on the superordinate-subordinate relation (hypernymy-hyponymy), which links more general synsets (e.g., “vehicle”) to increasingly specific ones (e.g., “car” and “sedan”).

This hierarchical structure is represented as a tree, and all noun hierarchies ultimately go up to the root node (“entity”). We used the Natural Language Toolkit (NLTK 3.4.5; https://www.nltk.org) to compute semantic similarity between prime and target according to three different methods (path length, Leacock-Chodorow, and Wu-Palmer), and the measure providing the highest correlation between priming predictions and the priming data (Wu-Palmer) was used in subsequent analyses.

We also evaluated two ad hoc models based on category membership. Unlike WordNet, these models computed semantic similarity based only on the categories and subcategories that appear in the stimulus set. *Taxonomic A* was coarser, consisting of 10 categories [subcategories]: Abstract [Mental State, Other Abstract], Event, Animate Object, Inanimate Object [Artifact, Food, Other Inanimate], and Place. *Taxonomic B* was more fine-grained, consisting of 21 categories [subcategories]: Abstract [Mental State, Other Abstract], Event [Social Event, Physical Event, Time Period], Animate Object [Human, Animal, Body Part], Inanimate Object [Artifact [Tool, Building Part, Furniture/Appliance, Instrument, Vehicle, Other Artifact], Food, Other Inanimate], and Place [Natural, Manmade].

### Distributional models

*Latent Semantic Analysis* (LSA; Landauer & Dumais, 1997) is one of the oldest and best-known techniques for generating word vector representations based on word co-occurrence statistics. Starting with a large set of text documents, LSA generates a matrix with unique words as rows and documents as columns, where each cell encodes the frequency with which a given word occurs in a given document. Since words with similar meanings tend to occur in similar contexts, the similarity between two rows is a measure of semantic similarity between the corresponding words. Singular value decomposition (SVD) is then used to extract independent factors (typically 300) from this matrix while preserving the similarity relations between words, resulting in a matrix of words by factors. The cosine similarity between these word vectors can predict human performance in semantic tasks (Landauer and Dumais, 1997, Pereira et al., 2016) and correlates with semantic similarity as measured by priming and as judged by human raters (Günther, Dudschig, & Kaup, 2016a, 2016b; Jones, Kintsch, & Mewhort, 2006). We generated LSA-based cosine similarity values from pre-trained 300-dimensional word vectors created from a two-billion word corpus (including Wikipedia, the British National Corpus, and the ukWaC corpus) divided into 5.4 million individual documents, obtained from the University of Tubingen’s repository for semantic spaces (http://www.lingexp.uni-tuebingen.de/z2/LSAspaces).

*Hyperspace Analogue to Language* (HAL) uses a words-by-words co-occurrence matrix, which is populated by counting word co-occurrences within a directional context window of a few words in each direction. The co-occurrences are weighted with the distance between the words, so that words that occur next to each other get the highest weight, and words that occur on opposite sides of the context window get the lowest weight. The result of this operation is a directional co-occurrence matrix in which the rows and the columns represent co-occurrence counts in different directions. We used HAL vectors created from a two-billion word corpus, which was created by concatenating the British National Corpus (BNC), the ukWaC corpus and a 2009 Wikipedia dump. They were also obtained from the University of Tubingen’s repository for semantic spaces. This space was built using a moving window with a size of 5 (2 to the left, 2 to the right), with the 100k most frequent words in the corpus as row words as well as content (column) words for the co-occurrence matrix.

Positive Pointwise Mutual Information weighting was applied, and Singular Value Decomposition was used to reduce the space from 100k to 300 dimensions.

*Word2vec* (Goldberg & Levy, 2014; Mikolov, Chen, Corrado, & Dean, 2013) is a distributional model that, rather than directly computing word co-occurrence frequencies, uses a 3-layer neural network trained to predict a word based on the words preceding and following it. We used the 300- dimensional word vectors trained on the Google News dataset (approximately 100 billion words) based on the continuous skipgram algorithm and distributed by Google (https://code.google.com/archive/p/word2vec). In contrast to word2vec, which uses only local information, *GloVe* (Pennington, Socher, & Manning, 2014) – short for Global Vectors – is based on the ratio of co-occurrence probabilities between pairs of words across the entire corpus. We used the 300-dimensional word vectors trained on Common Crawl (840 billion words) and made available by the authors (https://nlp.stanford.edu/projects/glove). In a comparative evaluation of distributional semantic models (Pereira, Gershman, Ritter, & Botvinick, 2016), word2vec and GloVe were the two top performing models in predicting human behavior across a variety of semantic tasks. Finally, fastText is a more recently developed model based on a continuous-bag-of-words (CBOW) algorithm that has been shown to outperform both word2vec and GloVe in standard benchmark tests of natural language processing (Mikolov, Grave, Bojanowski, Puhrsch, & Joulin, 2017). It contains several improvements over the original word2vec model (e.g., position-dependent weighting and use of subword information). We used the pre-trained vectors provided by the authors (https://fasttext.cc/docs/en/english-vectors.html), which were trained with subword information on Common Crawl (600 billion words).

### Participant-generated features

The *Semantic Feature Production Norms* (SFPN) devised by McRae and colleagues (2005) are arguably the largest and best-known effort to characterize word meanings in terms of semantic features. They have been used to test a variety of claims about the organization of the semantic system (e.g., Cree, McRae, & McNorgan, 1999; Cree and McRae, 2003; O’Connor, Cree, & McRae, 2009). Features are derived from descriptive properties generated by human participants in a property listing task. The properties are subsequently standardized into features with a binary value (present/absent), and their frequencies computed for each concept. This procedure results in vector-based concept representations based on thousands of features. The features represent various types of information, including perceptual (e.g., “is red”, “roars”), taxonomic (e.g., “is a mammal”), goal-related (e.g., “used for cooking”), and contextual association, as well as complex conceptual properties (e.g., “lays eggs”, “lives in the water”). We used the list of cosine similarities provided by Buchanan, Valentine, & Maxwell (2019), downloaded from https://github.com/doomlab/Word-Norms-2, to generate the SFPN model predictions.

### Control ratings-based model

To investigate the possibility that models based on human ratings of semantic features might have an inherent advantage over other kinds of models regardless of the kind of information they encode, we constructed a model based on a set of 12 features of word meaning that do not clearly map onto major, independently defined functional brain systems: “Familiarity”, “Musicality”, “Selfness”, “Number”, “Complexity”, “Approaching”, “Time”, “Communicative purpose”, “Large”, “Path”, “Dark”, and “Happy” (*Control12*; see **Supplemental Table 4** for details).

### Data Analysis

Analyses were conducted in Python using custom scripts. Only trials that received correct responses were included in the analyses. For each testing session, we computed the mean and standard deviation of response times (RTs) for correct trials. Trials with RTs more than 5 standard deviations away from the session mean for each participant were considered extreme values and excluded from further analyses.

RTs were standardized (z score) by testing session, separately for each participant. Priming was computed for each target as the difference in standardized RT between the two sessions. For all semantic models other than WordNet, semantic similarity was computed as the cosine between the vector representations of the two words. Priming predictions were computed for each model as P = similarity(Target, Prime B) – similarity(Target, Prime A) The correspondence between model predictions and observed priming (i.e., RSA correlation) was computed via Spearman correlations. Correlations were tested for significance across participants (subject-wise analysis; SW for short) using the Wilcoxon signed rank test, one-tailed. We also computed the Spearman correlations between model predictions and the group-averaged priming. This procedure averages out some of the noise in the data, resulting in much higher correlation scores. These correlations were tested using permutation tests (group-averaged priming analysis, or GA for short). We also conducted separate analyses for concrete and abstract target words to explore a possible effect of target concreteness on the results. Differences in correlation strength between predictions from different models were assessed via two-tailed signed rank test (SW) and Steiger’s Z test for dependent correlations (GA).

As an additional statistical approach, we conducted a set of linear regression (ordinary least squares) analyses of group-averaged priming including one regressor (i.e., predictions from a single semantic model) at a time and compared AIC across models. In another set of analyses, a linear regression model was fit, separately for each semantic model and for each participant, using the model’s priming predictions as the predictor variable, producing an AIC index for each. The AIC difference between two models was tested for significance across subjects with Wilcoxon’s signed rank tests (two-tailed).

The noise ceiling of a dataset corresponds to the highest RSA correlation that any model could achieve given the amount of noise in the data (Nili, Wingfield, Walther, Su, Marslen-Wilson, & Kriegeskorte, 2014). We computed the upper-bound noise ceiling estimate as the mean, across all participants, of the Spearman correlations between priming magnitude for each participant and the group mean. The lower-bound noise ceiling estimate was computed as the mean, across all participants, of the Spearman correlations between priming magnitude for each participant and the mean priming magnitude across all *other* participants.

Data reliability was also assessed via split-half correlations with 10,000 iterations. On each iteration, the subject sample was randomly split into 2 halves (15 and 16 subjects each) and the average item-level priming magnitudes were calculated for each half separately. The correlation between priming magnitudes of the two halves was computed and the Spearman–Brown formula was applied to the result. We report the average correlation across all iterations.

Partial correlations were used to evaluate the extent to which a model predicted the observed priming magnitudes once the predictions of a different model were taken into account (i.e., pairwise partial correlations). This provided a measure of how much unique information about word similarity patterns (as measured via priming) was encoded in each model. The main goal of these analyses was to verify whether each experiential model explained unique variance in the priming data beyond what could be explained by each distributional and taxonomic model. For that purpose, we computed the partial Spearman correlation between priming and the predictions of each experiential model (Exp48 or Lancaster9) after partialling out the predictions of each of the other (non-experiential) models, one at a time. We also computed the partial correlation between the predictions of each of the other models and priming after partialling out the predictions of each experiential model (one at a time) for comparison. We supplemented these analyses with ordinary least squares regression models including two regressors (predictions from two semantic models) for all pairs of models.

We also used partial correlations to assess each semantic model in terms of how much of the variance in the priming data it could uniquely explain while controlling for the other two kinds of information simultaneously. Thus, we computed the partial Spearman correlation between the predictions of each model and priming after partialling out the predictions of the best performing model of each of the other two kinds. For example, the correlation between priming and the predictions of each experiential model after partialling out the variance explained by the best distributional model (GloVe) and the best taxonomic model (Taxonomic B) simultaneously, denoted as Exp48[GloVe+TaxB] and Lancaster[GloVe+TaxB]. We did this for each semantic model (i.e., word2vec[Exp48+TaxB], Glove[Exp48+TaxB], fastText[Exp48+TaxB], TaxonomicA[Exp48+GloVe], and so on).

As additional metrics to help evaluate the performance of the semantic models, we also assessed the extent to which the subjective similarity and association ratings predicted semantic priming using Spearman correlation. We used partial correlation to verify whether each representational model predicted priming above chance level while controlling for each of these ratings, which would indicate that the kinds of information uniquely encoded in that model are more important for the neural representation of word meaning than suggested by explicit ratings alone.

## Results

### Lexical decision

Group mean accuracies (Acc) and response times (RTs) in the lexical decision task are presented in **Table 1**. The classic RT advantage for words over pseudowords was observed (p = 2 x 10^-9^, paired-samples t-test, two-tailed). RTs were also faster in session 2 compared to session 1 for both words and pseudowords (both p < .0004, paired-samples t-test, two-tailed), indicating that performance on the task improved with practice even though sessions were separated by several days. Importantly, the assignment of specific primes to the first or second session was counterbalanced across participants, so that the effect of session order was cancelled out in the computation of priming. Split-half reliability of the magnitude of the priming effect was .52, which is high for semantic priming in lexical decision (Heyman et al., 2018). The difference in prime-target orthographic Levenshtein distance between the two sessions did not correlate with priming (ρ = .00), neither did differences in the other 23 lexical variables listed in **Supplemental Table 2**, except for concreteness (ρ = .18, uncorrected p = .022), ruling out the possibility that priming magnitude was driven by ortho-phonological factors.

**Table 1.** Group mean accuracy (Acc) and response time (RT) in the lexical decision task. Number in parenthesis is the standard error of the mean.

|  | Session 1 |  | Session 2 |  | Overall |  |
| --- | --- | --- | --- | --- | --- | --- |
|  | Word | Pseudoword | Word | Pseudoword | Word | Pseudoword |
| Acc | 0.97 (.006) | 0.95 (.008) | 0.97 (.005) | 0.96 (.006) | 0.97 (.004) | 0.96 (.005) |
| RT (ms) | 631 (14) | 722 (20) | 598 (15) | 662 (19) | 614 (10) | 692 (14) |

### Semantic similarity rating

The median Pearson correlation between the ratings from each participant and the group median ratings was r = .92. For 20 of the 23 participants, correlations were between .86 and .96. The correlation was somewhat lower for one participant (.54). For two participants, correlations were exceptionally low (.11 and .26) indicating poor compliance with task instructions; these two participants were thus excluded from further analyses. Split-half correlation for the remaining 21 participants was .98, indicating excellent reliability.

### Thematic association rating

The median Pearson correlation between the ratings from each participant and the group median ratings was r = .86. Four participants were excluded due to exceptionally low correlation with the group median ratings (r < .06) indicating poor compliance with task instructions. For the remaining 21 participants, correlations ranged from .39 to .93. Split-half correlation was .94, again showing excellent reliability.

Correlations among model predictions are shown in **Supplemental Table 5**. As shown in **Figure 2, Supplemental Figure 2**, and **Supplemental Data 1**, all semantic representation models (including Control12) and relatedness ratings predicted priming significantly above chance, in subject-wise analyses (SW; all mean ρ > .10, all FDR-corrected p < 10^-6^, signed rank test) as well as in analyses based on group-averaged priming (GA; all ρ > .34, all FDR-corrected p < 10^-4^, permutation test) analyses. RSA correlations approached the lower-bound estimate of the noise ceiling (0.147) for all models except Control12, indicating that differences in performance between models may have been masked by a ceiling effect imposed by the level of noise present in the priming data. The same was true for the explicit ratings of subjective semantic similarity and thematic association. Except for LSA, all models predicted priming significantly more accurately than Control12 in the SW analysis (all mean differences in ρ > .02, all FDR-corrected p < .05, signed rank test, two-tailed). In the GA analysis, Exp48, GloVe, fastText, Taxonomic A, and Taxonomic B outperformed Control12 (all differences in ρ > .13, all FDR-corrected p < .03). The Exp48 model was the only one whose RSA correlation numerically surpassed the lower-bound noise ceiling estimate or the predictions derived from the semantic similarity ratings, and the only one whose correlation was stronger than that achieved by LSA in both SW and GA analyses (SW: FDR-corrected p = .045, signed rank test, two-tailed; GA: FDR-corrected p = .024, permutation test, two-tailed). In the GA analysis, Exp48 accounted for 27% of the variance in the data (Pearson’s r = .52). The overall pattern of model performances was similar across concrete and abstract targets (**Supplemental Figure 3**), although correlations for abstract targets had generally lower values and larger variances, possibly as a result of the smaller number of trials (53 abstract versus 157 concrete). In the ordinary least squares analysis of group-averaged priming including one regressor (i.e., predictions from a single semantic model), AIC values for each model produced a similar pattern of results (**Supplemental Tables 7** and **8).**

**Figure 2.**
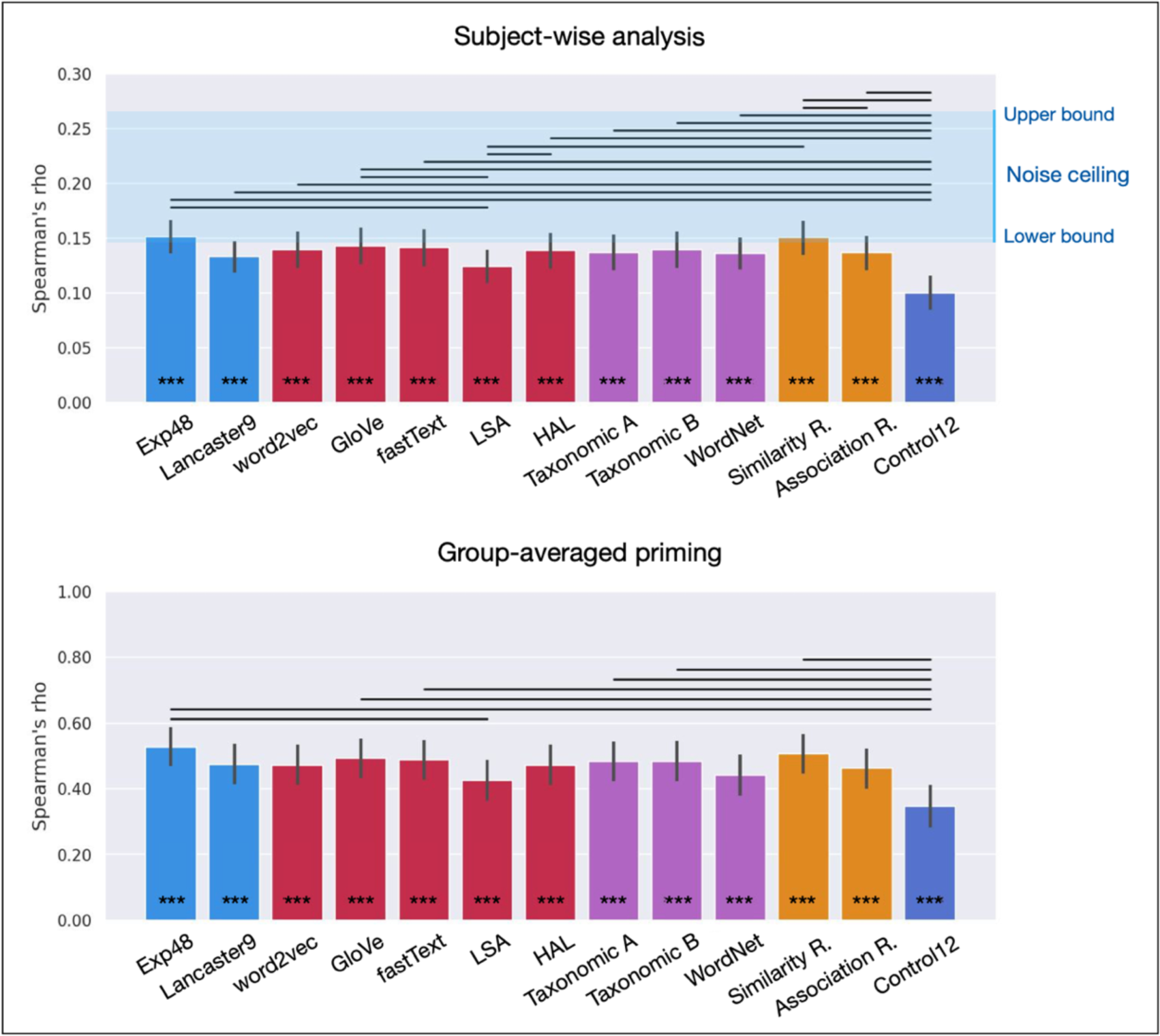
RSA correlations for experiential (light blue), distributional (red), and taxonomic (purple) models of semantic representation, for explicit ratings of semantic similarity and thematic association (orange), and for a control model based on a set of feature ratings that does not reflect the functional organization of the brain (dark blue). Horizontal lines indicate significant differences between correlations, FDR-corrected p < .05. *** FDR-corrected p < .0001.

Crucially, partial correlation analyses revealed that the experiential models explained unique variance in the priming data that could not be accounted for by taxonomic and distributional semantics models. Pairwise partial correlations evaluated the unique effect of each experiential model (blue bars in **Figure 3**, left-side panels) while controlling for each of the other models, one at a time, and, conversely, the unique effects of each taxonomic (purple) and distributional (red) model while controlling for each of the experiential models (see **Supplemental Data 2** and **3**). All partial correlations were highly significant for Exp48 (all FDR-corrected p < .002) while none of the taxonomic or distributional models correlated with priming when Exp48 was partialled out (all FDR-corrected p > .08). In other words, none of the other models explained significant variance beyond that explained by Exp48, while Exp48 explained significant unique variance not explained by any other models. This is strong evidence that experiential information is directly encoded in lexical concepts.

**Figure 3.**
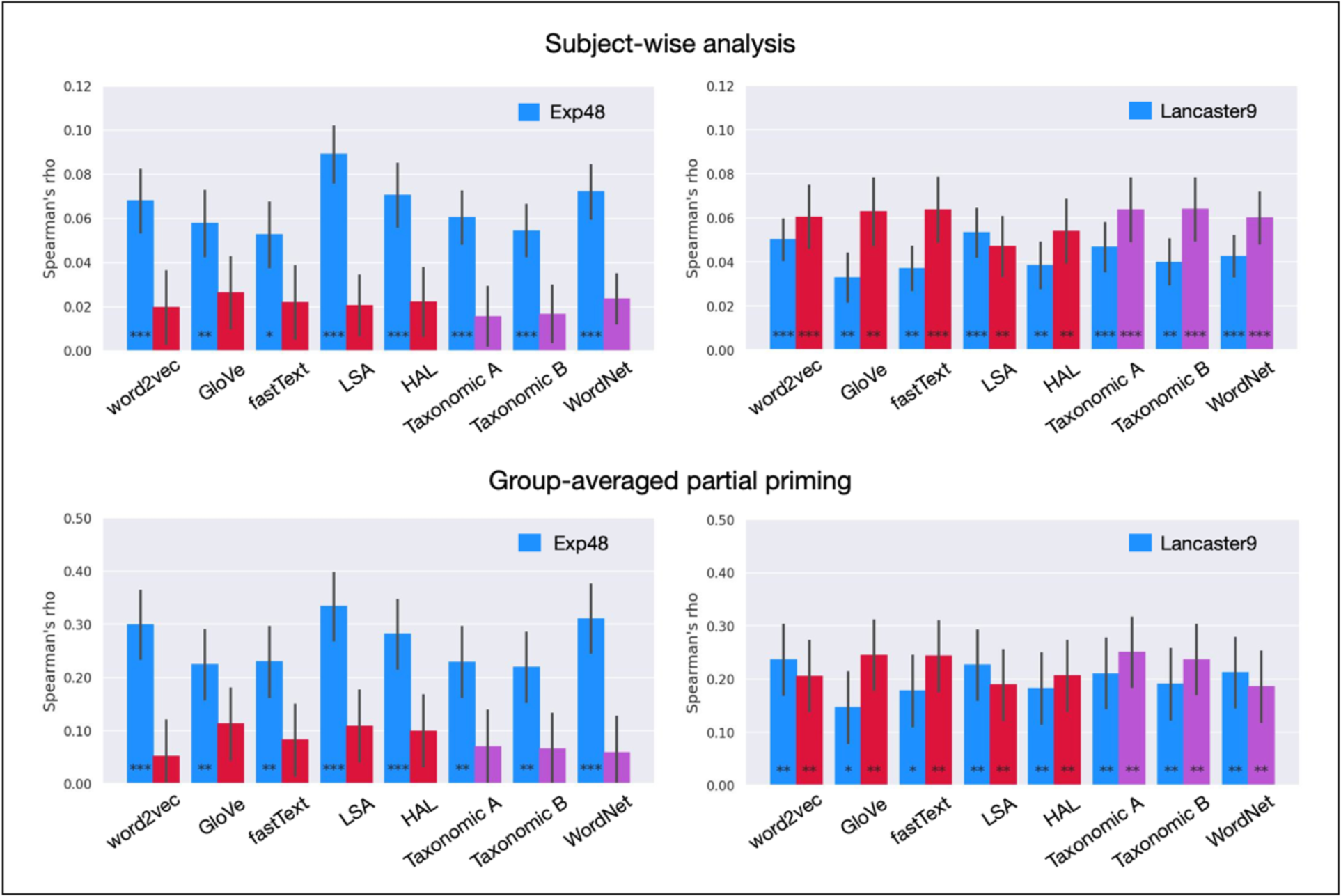
Pairwise partial correlations evaluating the unique variance explained by the experiential models (blue) while controlling for each of the other models, and the unique variance explained by the distributional (red) and taxonomic (purple) models while controlling for each experiential model. ***FDR- corrected p < .0005; ** < .005; * < .01.

Interestingly, given its relatively small number of features, the Lancaster9 sensory-motor model also explained unique variance in priming after controlling for any of the distributional, and taxonomic models (blue bars in **Figure 3**, right-side panels, and **Supplemental Data 3**). This indicates that information about the basic sensory-motor structure of the brain is encoded in concept representations and is automatically activated within 200 ms after word presentation.

The pairwise partial correlation for Exp48 controlling for Lancaster9 predictions (and vice versa) showed that Exp48 accounted for all the variance in priming explained by Lancaster9 plus additional unique variance (SW: Exp48 mean rho = .079, p = 2.7 x 10^-6^; Lancaster9 mean rho = .010, p = .24, signed rank test; GA: mean rho = .29, p = .0001; Lancaster9 mean rho = .07, p = .14, permutation test; Supplemental Data 5). This difference in partial correlations was significant (SW: p = .011, signed rank test; GA: p = 2.1 x 10^-7^, Steiger Z test; both two-tailed). Ordinary least squares models including two regressors (i.e., predictions from two semantic models), for all possible pairs of models, produced similar results (**Supplemental Table 9).**

Separate analyses for concrete and abstract targets suggested an overall advantage for the experiential models with both kinds of concepts, although not all partial correlations for those models reached significance, likely due to insufficient power (**Supplemental Figure 4**, blue bars).

Exp48 was the only model whose predictions correlated with priming when the variance explained by the best performing model of each of the other two kinds was partialled out (i.e., Exp48[GloVe+TaxB]; SW: mean ρ = .043, FDR-corrected p = .03, signed rank test; GA: ρ = .176, FDR-corrected p = .021, permutation test). The correlation was marginally significant for Lancaster9[GloVe+TaxB] (SW: mean ρ = .025, FDR-corrected p = .050; GA: ρ = .127, FDR-corrected p = .066) and not significant for any of the other models (SW: all mean ρ < .021, all FDR-corrected p > .262; GA: all mean ρ < .088, all FDR-corrected p > .418). Linear regression models including three regressors at a time showed that only Exp48 and Lancaster9 predicted priming significantly above chance when simultaneously controling for the predictions of the top-performing semantic model of each of the other two types (**Supplemental Table 10)**.

Exp48 predicted priming above chance level even after partialling out the variance explained by all taxonomic and distributional models simultaneously (SW: mean ρ = .039, FDR-corrected p = .021, signed rank test; GA: ρ = .188, FDR-corrected p = .012, permutation test), again providing strong evidence that experiential information is independently encoded in lexical semantic representations and plays a central role in semantic word processing (**Supplemental Figure 5**). Lancaster9 predicted priming after simultaneously partialling out the variance explained by all five distributional models (SW: mean ρ = .0293, FDR-corrected p = .019, signed rank test; GA: ρ = .153, FDR-corrected p = .027, permutation test) or by all three taxonomic models simultaneously (SW: mean ρ = .029, FDR-corrected p = .006, signed rank test; GA: ρ = .150, FDR-corrected p = .030, permutation test), although correlations were only marginally significant when all eight models were partialled out simultaneously (SW: mean ρ = .0184, FDR-corrected p = .067, signed rank test; GA: ρ = .124, FDR-corrected p = .074, permutation test).

We then assessed whether each semantic model predicted priming above chance level after partialling out the variance explained by the subjective semantic similarity ratings. Except for Exp48 (GA: FDR p < .05, permutation test), no other models showed significant partial correlations after correcting for multiple comparisons. Only Exp48, Lancaster9, and Taxonomic B reached uncorrected significance (SW: p = .007, p = .014, and p = .030, respectively, signed rank test; GA: p = .003, p = .027, and p = .046, respectively, permutation test; **Supplemental Figure 6**, left-hand panels). These results suggest that subjective ratings of semantic similarity do not completely capture the importance of experiential information for semantic priming. Partial correlations did not approach significance for any of the distributional models (all uncorrected p > .28) nor for Control12 (both SW and GA p > .96). All experiential, distributional, and taxonomic models predicted priming after partialling out the thematic association ratings (all FDR-corrected p < .034; **Supplemental Figure 6**, right-side panels), indicating that their performance in the main analysis was primarily driven by semantic similarity. Control12 was the only model whose predictions did not correlate with priming when controlling for the association ratings (both SW and GA p > .58), in stark contrast to the experiential models.

To evaluate whether the good performance of the experiential models in predicting priming was driven by the information source (i.e., explicit ratings of the importance of semantic features to word meaning) rather than by their information content (about the sensory-motor-affective organization of experience), we conducted pairwise partial correlations comparing Control12 to each of the other models (**Supplemental Data 6**). **Figure 4** shows that Control12 did not perform nearly as well as Lancaster9 did against distributional and taxonomic models (compare to **Figure 3**, right side panels), despite containing a larger number of features. It also shows that most of the variance explained by Control12 was also explained by Lancaster9, but the reverse was not true. This confirms that model performance was driven primarily by representational structure, not by the type of behavior from which model values were derived.

**Figure 4.**
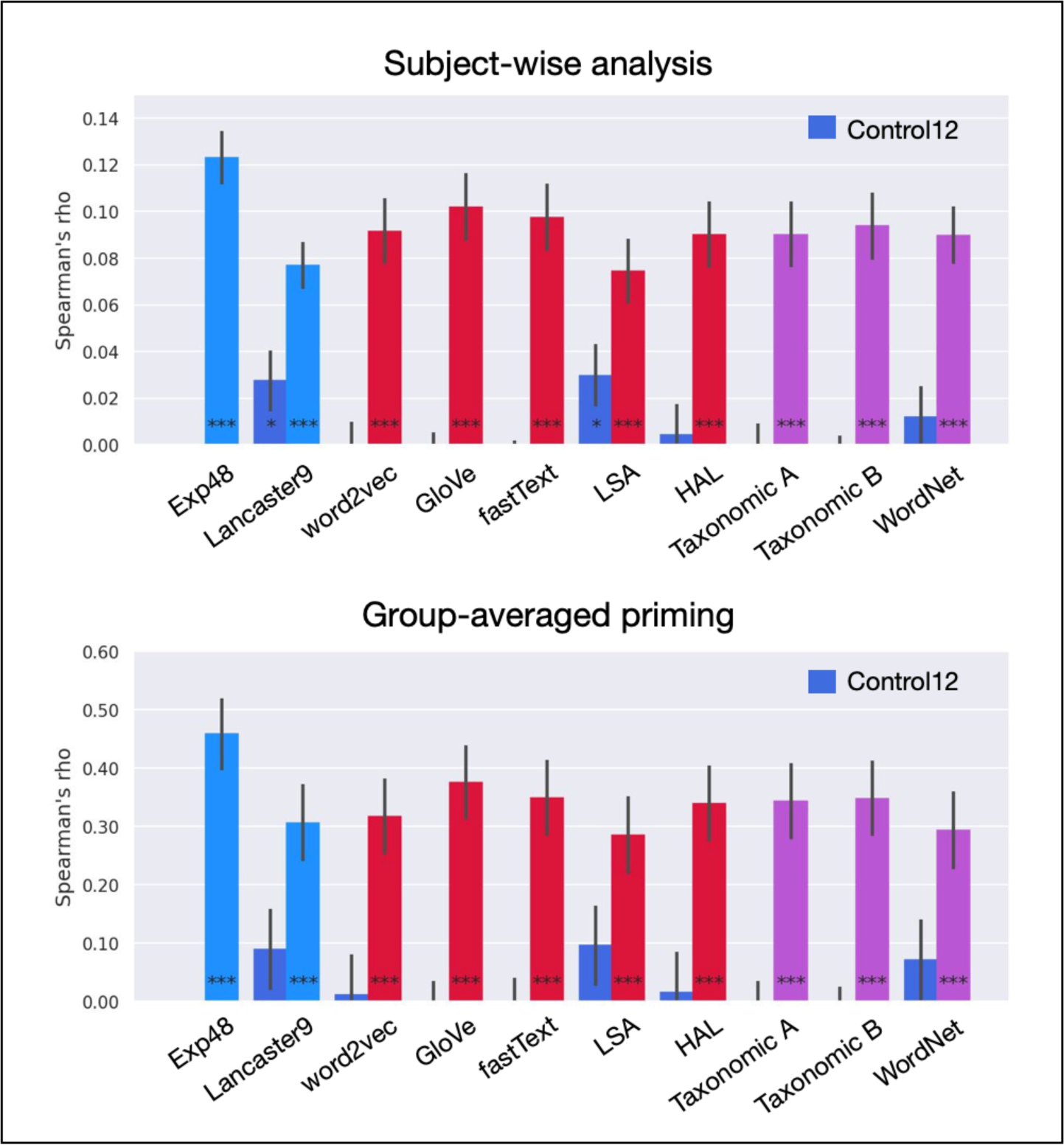
Pairwise partial correlations evaluating the unique variance explained by the Control12 model (dark blue) while controlling for the variance explained by each of the 10 semantic models evaluated, and vice versa. ***FDR-corrected p < .0005; ** < .005; * < .05.

The RSA including the SFPN model, including only trials for which those norms were available, showed that this model also succeeded in predicting priming magnitude (SW: mean ρ = .145, FDR-corrected p = 1.4 x 10^-7^, signed rank test; GA: ρ = .458, FDR-corrected p = .0001, permutation test; **Figure 5 and Supplemental Data 4**). Results for the other models were similar to those in the main analysis. There were no differences in correlation strength between SFPN predictions and those from other models (all FDR-corrected p > .27). However, pairwise partial correlations showed that, overall, it performed worse than the other three kinds of models (**Figure 6** and **Supplemental Data 4)**. Partial correlations for this model’s predictions did not reach significance when partialling out the variance explained by either GloVe, fastText, WordNet, or Exp48 (SW and GA: all FDR-corrected p > .05, signed rank and permutation tests), but they were significant for all other models when controlling for the effect of SFPN (SW: all p < .008, signed rank test; GA: all p < .027, permutation test), except in the GA analysis for LSA (p = .085, permutation test). This finding suggests that semantic features spontaneously produced by study participants with the goal of characterizing word meaning are worse predictors of automatic semantic priming than the other kinds of models investigated.

**Figure 5.**
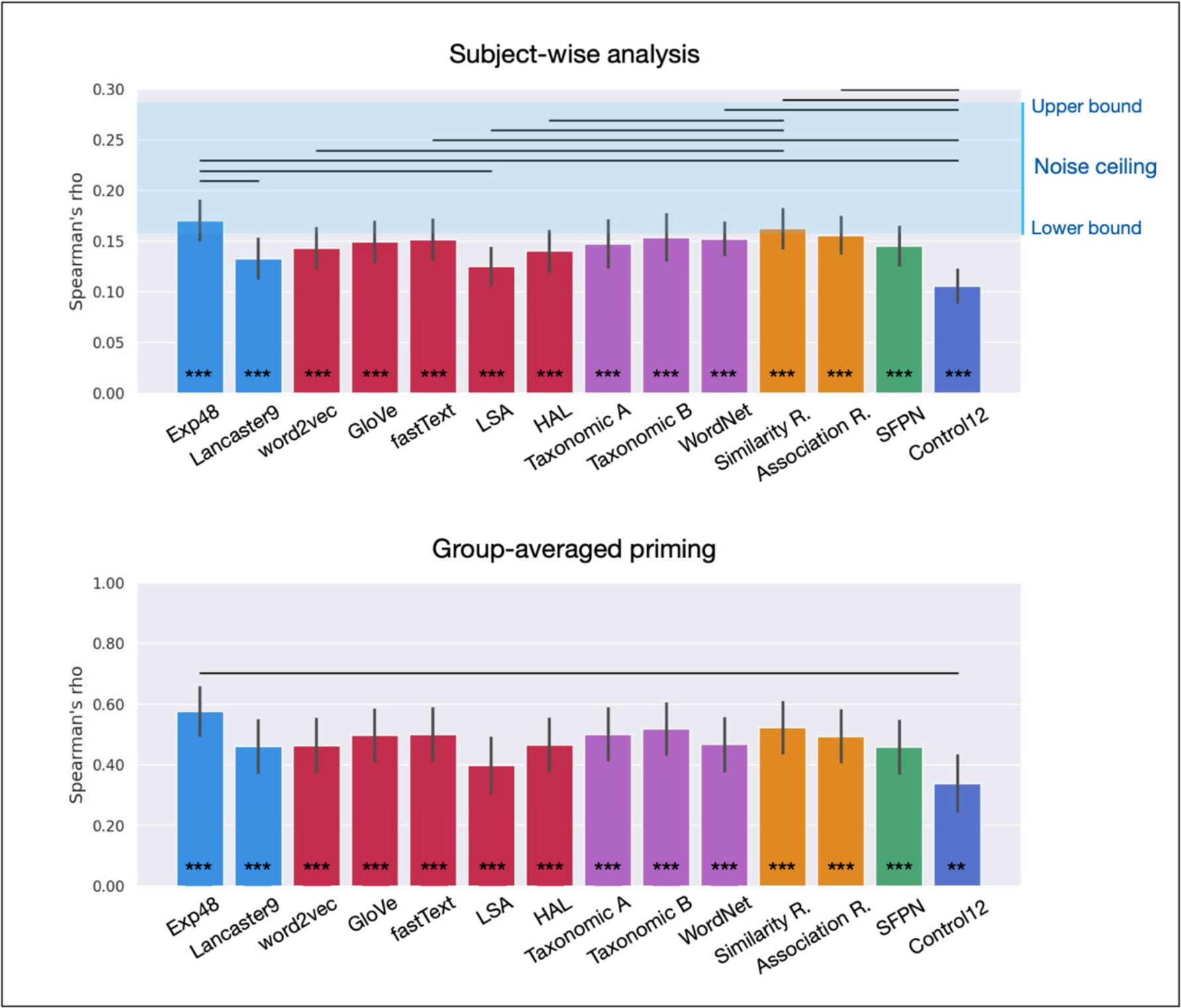
RSA results including only the 97 trials for which SFPN scores are available. Horizontal lines indicate significant differences between RSA correlations, FDR-corrected p < .05. ***FDR-corrected p < .0005; ** < .005; * < .05.

**Figure 6.**
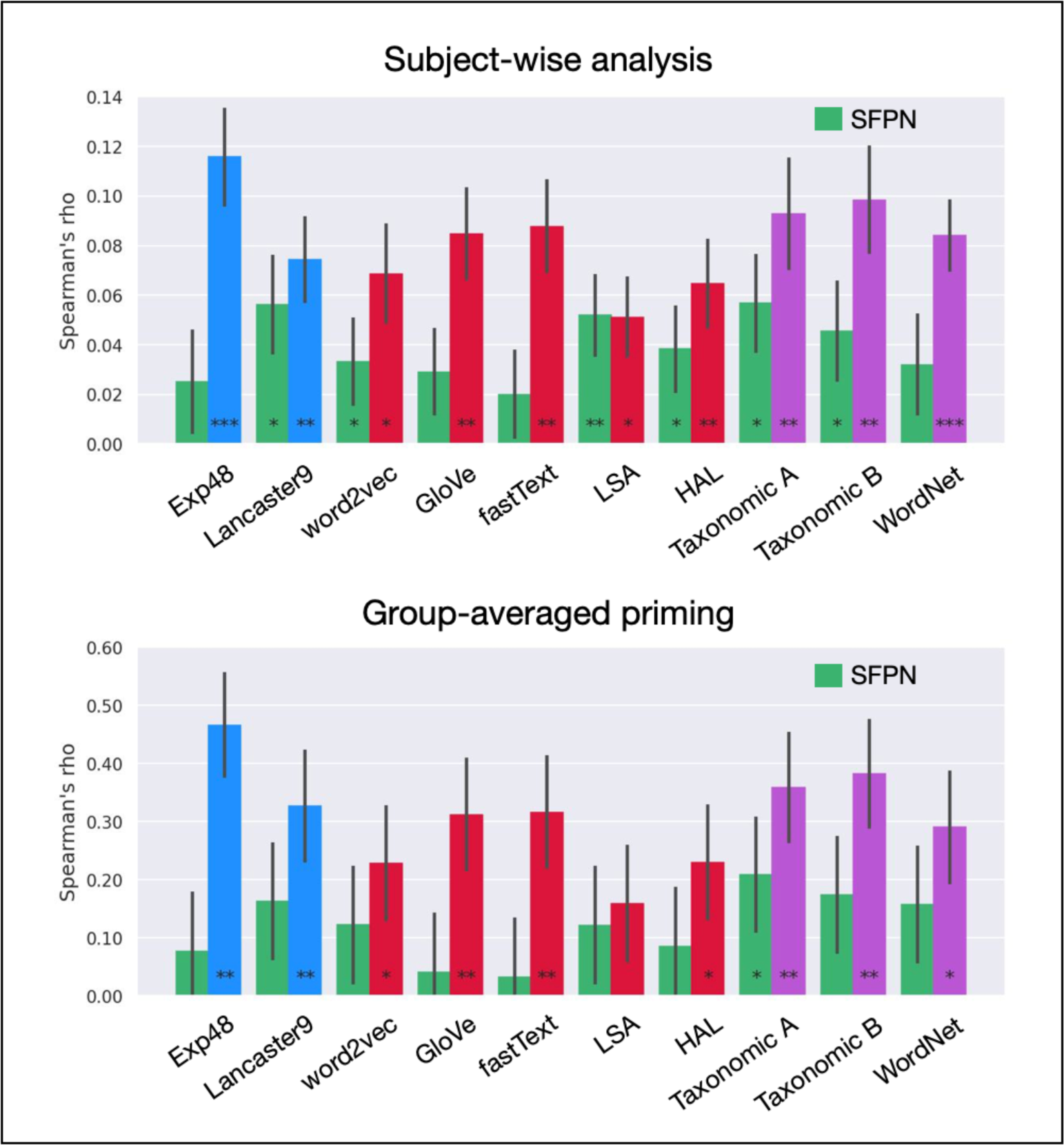
Pairwise partial correlations evaluating the unique variance explained by the SFPN model (green) while controlling for predictions of each of the other models, and the unique variance explained by the other models while controlling for the SFPN predictions. ***FDR-corrected p < .0005; ** < .005; * < .05. (includes only the 97 trials for which SFPN scores are available).

Finally, we evaluated the extent to which priming was driven by semantic similarity relative to contextual association between prime and target. The partial correlation between semantic similarity and priming while controlling for contextual association was significant (SW: ρ = 0.06, p = 0.0001, signed-rank test; GA: ρ = 0.21, p = 0.003, permutation test), while the partial correlation for contextual association while controlling for semantic similarity was not (SW: ρ = -0.002, p = 0.78, signed-rank test; GA: ρ = -0.021, p = 0.76, permutation test). This indicates that priming was primarily driven by feature-based similarity, with little (if any) contribution of contextual association relationships.

## Discussion

Automatic semantic priming is widely regarded as an important source of information about how word meanings are represented and accessed during language processing (e.g., Burgess & Lund, 2000; Giffard, Desgranges, Nore-Mary, Laleveé, Sayette, et al., 2001; McRae, de Sa, Seidenberg, 1997; Mandera et al., 2017). Psycholinguistic research indicates that its magnitude can reflect various types of semantic relations between words, including thematic association, feature-based similarity, taxonomic relationships, and frequency of word co-occurrence in natural language use (Chiarello and Richards, 1992; Landauer & Dumais, 1997; Pereira, Mandera et al., 2017; McRae, Sa, & Seidenberg, 1997; Perea & Rosa, 2002; Wingfield & Connell, 2022). We set out to evaluate the extent to which automatic semantic priming reflects three distinct types of information, namely experiential, distributional, and taxonomic similarity. While previous studies had investigated whether particular sensory-motor or affective features play a role in semantic word processing, the degree to which experiential information explains semantic behavior was still unknown. Using both RSA and linear model approaches, we assessed the combined effect of several experiential dimensions on the similarity structure of lexical concepts as revealed by automatic semantic priming. We evaluated two experiential models, one based on coarse dimensions (Lancaster) and one based on both coarse and fine-grained dimensions (Exp48), relative to five of the best known distributional models and to three taxonomic models. The results showed a remarkably robust association between priming and experiential similarity, with both experiential models accounting for unique variance in the data after regressing out the variance explained by distributional and taxonomic models (**Figure 3 and Supplemental Figure 4**) and by subjective ratings of semantic similarity and thematic association (**Supplemental Figure 6**). These results were found in subject-wise analyses as well as in analyses of group-averaged priming. Furthermore, they could not be explained by ortho-phonological factors (since none of those factors was correlated with priming) nor by grammatical class (since all the stimuli were nouns).

The finding that Exp48 accounted for all the variance in priming explained by Lancaster9 plus additional unique variance is consistent with the idea that lexical concepts encode information about the more fine-grained experiential features included in Exp48 but not in Lancaster9, and consistent with the results of Fernandino et al., 2022, where the equivalent effect was found for similarities between word-specific fMRI activation patterns. While experiential structure could still be detected at a coarse level (i.e., the relative importance of information originating from each of the major sensory-motor modalities, as indexed by the Lancaster9 model), information about the relative importance of more specific systems, such as those underlying the processing of color, shape, texture, emotional valence, and reward value, among others, seems to play a role in semantic word processing.

Much of the literature on embodied semantics has focused on whether modality-specific brain systems play a role in the representation of word meaning. However, it is important to note that seemingly elementary features of sensory-motor experience actually integrate information across multiple sensory-motor systems. The sense of space, for example, is a representational system integrating inputs from the visual, motor, tactile, proprioceptive, vestibular, and auditory systems, and is subserved by a network of multimodal neural structures such as the hippocampus, the entorhinal cortex, and the posterior parietal cortex (Baumann & Mattingley, 2014; Graziano, Hu, & Gross, 1997; Graziano, Yap, & Gross, 1994; Sanders, Rennó-Costa, Idiart, & Lisman, 2015).

Graspability and manipulability also integrate information across visual, tactile, proprioceptive, and motor systems, and appear to rely most strongly on multimodal cortical areas such as the posterior middle temporal gyrus, the anterior supramarginal gyrus, the anterior intraparietal sulcus, and the ventral precentral sulcus (Jastorff, Begliomini, Fabbri-Destro, Rizzolatti, & Orban, 2010; Peeters, Simone, Nelissen, Fabbri-Destro, Vanduffel, et al., 2009; Reynaud, Navarro, Lesourd, & Osiurak, 2019). The sense of causality, another ubiquitous aspect of human experience, is still poorly understood from a neurobiological perspective, but it is likely built upon experiential primitives that integrate sensory, motor, spatial, and temporal information into causal event schemas (Leshinskaya, Bajaj, & Thompson-Schill, 2021; Pelt, Heil, Kwisthout, Ondobaka, Rooij, et al., 2016; Pulvermüller, 2018; Rakison & Krogh, 2012). Therefore, while modality-specific effects on semantic language processing have provided evidence that sensory-motor systems contribute to semantic representation, many (if not most) sensory-motor features of experience combine information from multiple modalities.

Exp48 includes experiential features at several levels of coarseness and modality specificity. Coarse modality-specific features include Vision, Audition, Touch, Taste, and Smell; fine-grained modality-specific features include Color, Texture, Sound Pitch, and Temperature. Multimodal features consist of spatial, temporal, and causal dimensions considered fundamental for the organization of experience, such as Peripersonal Space (i.e., spatial proximity to the body, within reaching distance), Manipulability, Caused (i.e., causally affected), and Consequential (i.e., causal agent). Shape, although listed as a visual feature in Binder et al. (2016), most likely results from the integration of visual, tactile, and motor information (at least in neurologically typical individuals).

Although the present data do not allow us to specify the extent to which each of the features included in the Exp48 and Lancaster9 models affects semantic priming, they suggest that at least some of the more fine-grained and multimodal features in Exp48 contribute to the phenomenon in a substantial way.

We also found that Exp48 accounted for all of the variance that was explained by any of the distributional and taxonomic models (**Figure 3**, left panels). In other words, although the non-experiential model predictions were correlated with priming magnitude when assessed in isolation (**Figure 2**), partial correlation analyses revealed that distributional and taxonomic similarity only predicted priming to the extent that they predicted experiential similarity. Therefore, we found no evidence for the hypothesis that distributional information contributes to the representation of lexical concepts. This finding, however, may be related to technical limitations of the particular distributional models tested; it remains possible that a different variety of distributional model could predict unique priming variance after accounting for the similarity structure predicted by Exp48. On the other hand, it is important to note that while the Exp48 and Lancaster9 models are based exclusively on theoretical principles, with no consideration of decoding or predictive performance, the architectures and parameters of some of the distributional models evaluated here (word2vec, GloVe, and fastText) are driven by performance optimization in word prediction via supervised learning (Mikolov et al., 2013, 2017; Pennington et al., 2014). It stands to reason that an experiential model optimized for semantic decoding (for example, via optimization of feature weights) would perform substantially more accurately than Exp48 and, potentially, more accurately than any possible distributional model.

It has been proposed that distributional information may play a more important role in the representation of abstract concepts than in the representation of concrete concepts . Although most target words in the present study (157 words) were relatively concrete (median concreteness = 4.9, IQR = 0.2), 51 target words were relatively abstract (median concreteness = 2.5, IQR = 1.4), such as “role”, “motive”, and “fate”. When trials with concrete and abstract targets were analyzed separately, there was no indication that distributional models performed better on the latter, whether in absolute terms or relative to the performance of the other models (**Supplemental Figure 3**). In fact, partial correlations showed that only Exp48 and Lancaster9 explained significant unique variance for abstract targets, providing evidence against that proposal (**Supplemental Figure 4**). Of course, given the relatively low number of abstract trials, these results should be interpreted with caution.

Partialling out the predictions of the semantic similarity ratings or the contextual association ratings indicated that, for all models, RSA correlations with priming were primarily driven by semantic similarity (**Supplemental Figure 6**). The two relatedness ratings were strongly correlated with each other (r = .90), but partial correlations showed that only one – semantic similarity – accounted for unique variance in the priming data that could not be explained by the other. It is worth noting that, although these ratings reflect people’s intuitions about how related two words are, there is no reason to believe that naïve intuition perfectly reflects semantic similarity (or association) as encoded in the brain. A clear lesson from experimental psychology is that our intuitions are poor guides to how the mind actually works. Therefore, the semantic similarity ratings should be considered as one measure of semantic similarity, not as the gold standard. Interestingly, the finding that experiential models predicted semantic priming while controlling for the similarity ratings indicates that the semantic factors underlying this phenomenon are better explained by experiential feature ratings than by explicit semantic similarity ratings. This suggests that experiential features play a more important role in concept representation than what could be inferred based on similarity ratings alone.

Previous studies have indicated that sensory-motor features of lexical concepts can be activated to a variable extent during semantic language processing, depending on the demands of the task (Dam, Brazil, Bekkering, & Rueschemeyer, 2014; Goh, Yap, Lau, Ng, & Tan, 2016; Raposo, Moss, Stamatakis, & Tyler, 2009; Tousignant & Pexman, 2012). These findings raise the question of whether experiential features can be considered an essential aspect of lexical concepts rather than mere “post-semantic” representations recruited by task-related, strategic mechanisms (Weiskopf, 2010). Support for the former was provided by an event-related potentials (ERP) study by Muraki, Sidhu, & Pexman (2019), which suggests that experiential information is initially activated (within 200 ms of word onset) regardless of task-related factors and subsequently modulated by top-down processes according to task demands. The present study provides additional evidence for this view by showing that experiential features are quickly and reliably activated during lexical decision, a task that makes no explicit demands on semantic processing. While semantic content is implicitly activated and affects response latencies, lexical decision does not favor any particular kind of semantic information associated with the word form, providing an unbiased assessment of the information content of lexical concepts. Furthermore, prime and target words were presented in a backward-masked, short SOA procedure that emphasizes the effect of semantic features that are activated within 200 ms of the prime onset. Therefore, our results indicate that experiential features are automatically activated during semantic word processing.

The idea that lexical concepts possess a “core” set of features that are automatically activated regardless of the context has been called into question by Lebois, Wilson-Mendehall, and Barsalou (2014). They reviewed several studies demonstrating that a variety of features considered central to the meanings of words can be modulated by contextual factors such as task demands and the composition of the stimulus set. The authors concluded that semantic feature activation is always dynamic and context-dependent, and that distinctions between core and peripheral features are therefore meaningless. Although we agree with Lebois et al. that context-independent automaticity should not be considered the gold standard of semantic processing and that context-dependent features are not irrelevant, we disagree with their dichotomous characterization of automaticity and context-dependency. The reason is that any phenomenon that is typically considered automatic is also susceptible to modulation by contextual factors. Consider, for example, the patellar reflex: the leg extension is an automatic response to the mechanical stimulus delivered to the patellar tendon, yet the latency and magnitude of the response can be modulated by a variety of contextual factors. To say that a particular semantic feature is automatically activated by a word form means that, under normal conditions, activation of the orthographic or phonological word form representation is a sufficient condition for the activation of that feature; it does not mean that the degree to which the feature is activated cannot be modulated by contextual factors. Furthermore, different semantic features are not all equally susceptible to task-related factors. It is clear that, among the semantic features that can be associated with a particular word form, there are some that appear to be more consistently activated across different contexts (and across speakers of the language) than others.

Hence, we do see a meaningful distinction between features that show relatively little sensitivity to contextual factors and those that are strongly context-dependent. Because the activation of semantic representations in our paradigm – particularly those associated with the prime – is completely incidental to the task and occurs at a relatively short time scale, our results suggest that experiential features are fundamental constituents of word meaning.

We also examined whether semantic models based on human-generated ratings – such as Exp48 and Lancaster9 – have an inherent advantage over distributional models, which are based on information extracted from language corpora without input from human raters. Afterall, it is no surprise that the subjective rating of semantic similarity performed better than the distributional models in predicting semantic priming, since it is an explicit measure of how similar the brain perceives two words to be, while the similarity metrics produced by models such as LSA and word2vec are indirectly derived from word co-occurrence patterns. This issue can be analyzed in terms of whether human-generated semantic ratings necessarily reflect semantic similarity and whether they are necessarily better predictors of semantic priming than distributional model dimensions. It seems clear that, unlike ratings of semantic similarity between two words, rating of the importance of particular features to the meaning of a word are not direct measures of semantic similarity. The set of semantic features that participants are asked to rate is chosen by the experimenter and constitute a hypothesis about which features are important for determining how similar two words are. If the features rated are not particularly salient to how the brain encodes word meanings, they will not predict any measures of semantic similarity (i.e., neither similarity ratings nor semantic priming). Indeed, we found that all experiential, distributional and taxonomic models evaluated – including Lancaster9, consisting of only 9 sensory-motor dimensions – predicted priming more accurately than Control12, consisting of 12 semantic feature ratings that are not directly related to the sensory-motor structure of the brain. Moreover, GloVe and fastText (as well as Exp48 and WordNet) accounted for all the variance explained by the SFPN model (based on human-generated semantic features of word meaning) plus additional variance in the priming data, demonstrating that human-generated semantic features do not necessarily predict priming more accurately than distributional models.

Another important consideration relates to how the different kinds of semantic models evaluated in the present study compare in terms of explanatory value. Different distributional models emphasize distinct aspects of the statistical regularities with which words co-occur with one another during natural language use, and can reveal which of these aspects, if any, are relevant for the representational code used by the brain to encode meaning. They cannot, however, provide information about how this representational code relates to the codes involved in perceptual, motor, interoceptive, and valuative processes that characterize word referents, which is an essential step toward a scientific understanding of how lexical concepts come to refer to real-world objects, situations and events. Therefore, these models are epistemically disconnected from the learning processes involved in the acquisition of word meanings in young children, which are largely based on sensory-motor information about word referents (e.g., Landau, Smith, & Jones, 1988; Mandler, 2004; Peters & Borovsky, 2019; Singleton & Saks, 2023; Wojcik & Saffran, 2013). Distributional models indirectly encode experiential information because the patterns in which words co-occur in natural language are influenced by the co-occurrence patterns of their referents in the world.

However, this statistical correspondence can only carry a fraction of the experiential information available to the brain during concept acquisition, which is further diluted among several other factors affecting word co-occurrence patterns such as polysemy and cultural conventions. Furthermore, distributional models are completely blind to aspects of experience that are so fundamental to our daily existence that they are automatically factored into our communication (i.e., assumed) without being encoded into words (e.g., the fact that two solid objects typically cannot occupy the same location at the same time; the fact that we are almost always positioned on essentially flat surfaces to which all objects are always attracted; and so on). To put it another way, the words we produce are meant to be interpreted according to a shared, non-verbal model of the world. While distributional models can perform surprisingly well in some well-specified, language-based semantic tasks, these successes do not constitute evidence that these models encode human-like semantic representations until they are able to translate between words and their real-world referents as well as humans do.

Taxonomic models provide information about the kinds of things that are usually perceived as relatively similar or dissimilar by the brain. To the extent that they may explain semantic data beyond what is explained by distributional and experiential models, they would indicate that a different kind of information reflecting categorical distinctions – not one emerging from sensory-motor-affective experience nor from word co-occurrence – plays a significant role in concept representation.

Although taxonomic models do not specify what kind of information that would be, semantic categories could reflect different mechanisms of neural representation, different neuroanatomical locations, or different patterns of connectivity of the underlying cortical areas (Mahon & Caramazza, 2011). By itself, however, a priori taxonomic information cannot account for semantic distinctions between individual members of a category or for the flexibility with which people categorize entities in different contexts (Barsalou, 1983; Lakoff, 1987).

Experiential feature models directly relate concept representations to sensory-motor, affective, spatial, temporal, and other independently defined cognitive and neural domains. Therefore, unlike the individual dimensions of distributional models, experiential dimensions are directly interpretable and have explanatory value. Different models reflect different combinations of experiential features, which can inform about the role played by different modality-specific and multimodal neural systems in concept acquisition and representation. Therefore, these models can clarify the nature of the relationships between word meanings, their biological substrates, and their real-world referents. In principle, they can explain both categorical and distributional phenomena, that is, how and why concepts cluster together into categories and how they co-occur during language use.

In conclusion, the results reported here are consistent with a hierarchical model of word meaning in which lexical concepts consist of fuzzy sets of experiential features encoded at various levels of representation throughout the cortex (Fernandino, Binder, Desai, Pendl, Humphries, et al., 2016). At the bottom of the hierarchy, primary sensory-motor, interoceptive, and limbic/paralimbic areas encode simple modality-specific features. At the next level, specialized networks encode more elaborate features (e.g., 3D shape, spatial relationships, manipulability, edibility, sequential structure, causal relationships, etc.) based on information extracted from lower-level areas and processed according to innate connectivity constraints and ecological/pragmatic needs. At the top level, a set of heteromodal cortical hubs integrate information across those networks to encode complex, flexible assemblages of experiential features that can become associated with an arbitrary word form (i.e., a phonological/orthographic representation). During semantic word processing, the word form representation initially activates a set of high-level experiential features stored in the heteromodal hubs, which, in turn, activate lower-level features stored in other multimodal areas in a context-dependent manner. These lower-level features can, in turn, activate simpler features stored in early sensory-motor and affective areas depending on task demands.

We identify these heteromodal hubs with the angular gyrus, the anterior lateral temporal cortex, the parahippocampal cortex, the precuneus/posterior cingulate cortex, and the medial prefrontal cortex. Collectively known as the “default mode network” (DMN), these areas have been shown to be the ones that are most distantly connected to primary sensory and motor areas, and they appear to integrate information across all modalities (Fernandino, Binder, et al., 2016; Fernandino, Tong, Conant, Humphries, & Binder, 2022; Margulies, Ghosh, Goulas, Falkiewicz, Huntenburg, et al., 2016; Sepulcre, Sabuncu, Yeo, Liu, & Johnson, 2012; Tong, Binder, Humphries, Mazurchuk, Conant, et al., 2022). They are also highly interconnected with each other (Buckner, Andrews-Hanna, & Schacter, 2008) and have been shown to be jointly activated during semantic language tasks (Binder, Frost, Hammeke, Bellgowan, Rao, et al., 1999; Binder, Desai, Graves, & Conant, 2009; Wirth, Jann, Dierks, Federspiel, Wiest, et al., 2011) and during other tasks that depend primarily on information retrieved from long-term memory, such as remembering the past, thinking about the future, and daydreaming (Christoff, Gordon, Smallwood, Smith, & Schooler, 2009; Mason, Norton, Horn, Wegner, Grafton et al., 2007; McKiernan, Kaufman, Kucera-Thompson, & Binder, 2003; Spreng & Grady, 2010). We had previously shown that individual words could be decoded from fMRI activation patterns in these areas using a multimodal sensory-motor model, but not with a model based on ortho-phonological features of the corresponding word forms (Fernandino, Humphries, et al., 2016). Furthermore, the similarity structure of fMRI activation patterns in these regions predicts the semantic similarity structure of both object and event nouns, and it does so significantly more accurately when semantic similarity is estimated based on experiential features than when it is based on taxonomic or distributional information (Fernandino et al., 2022; Tong et al., 2022). In agreement with this model, the present study provides robust behavioral evidence that multimodal experiential information is automatically activated during semantic word processing.

Further research is required to determine in more detail the degree to which different experiential features contribute to the representation of lexical concepts and how their activation is modulated by contextual factors.

### Data availability

All data and computer code used to generate the results and figures are available at the Open Science Framework: https://osf.io/yp4mz/?view_only=ee24c3ab443a4240a93857e1fefe3565. DOI 10.17605/OSF.IO/YP4MZ

## Acknowledgements

The authors would like to thank Taylor Kalmus for help with data collection, Jeffrey Binder for helpful discussions and invaluable support, and two anonymous reviewers for helpful comments and suggestions on a previous version of this manuscript. This work was supported by an NIDCD grant R01DC016622.

## Supplemental figures

**Supplemental Figure 1.**
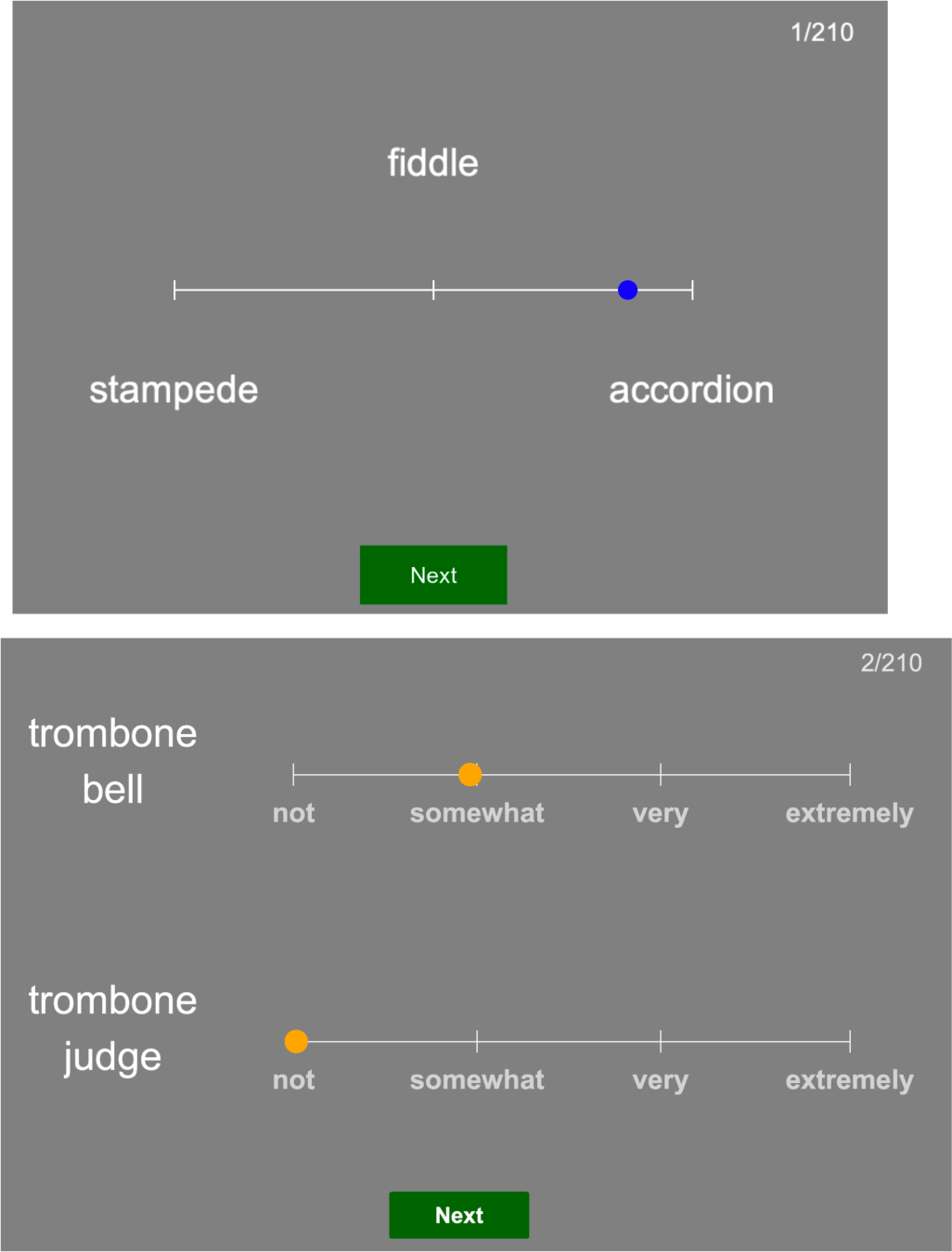
Examples of the semantic similarity (left) and thematic association (right) rating tasks.

**Supplemental Figure 2.**
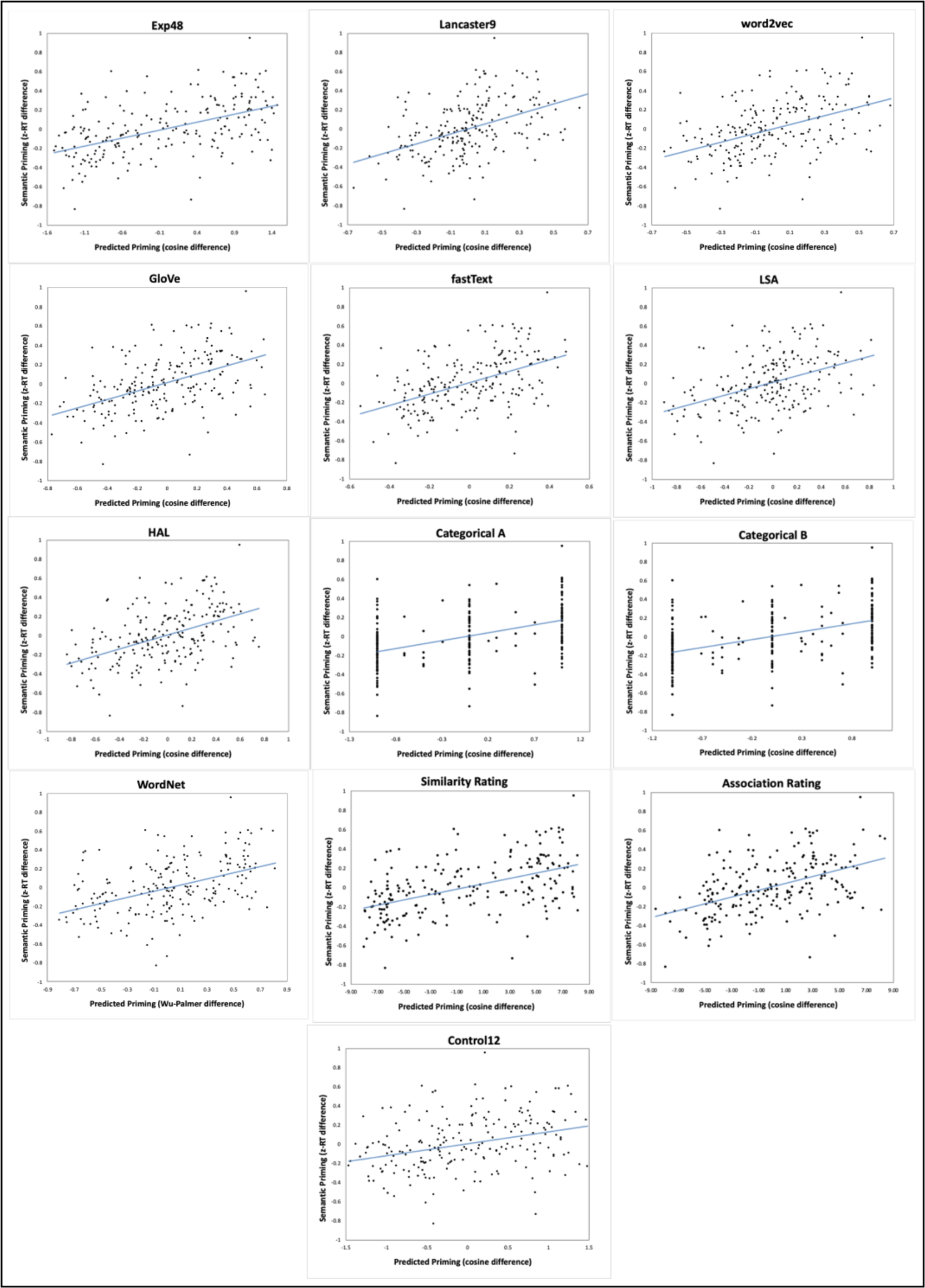
Scatterplots of semantic priming by predicted priming for each model and relatedness rating.

**Supplemental Figure 3.**
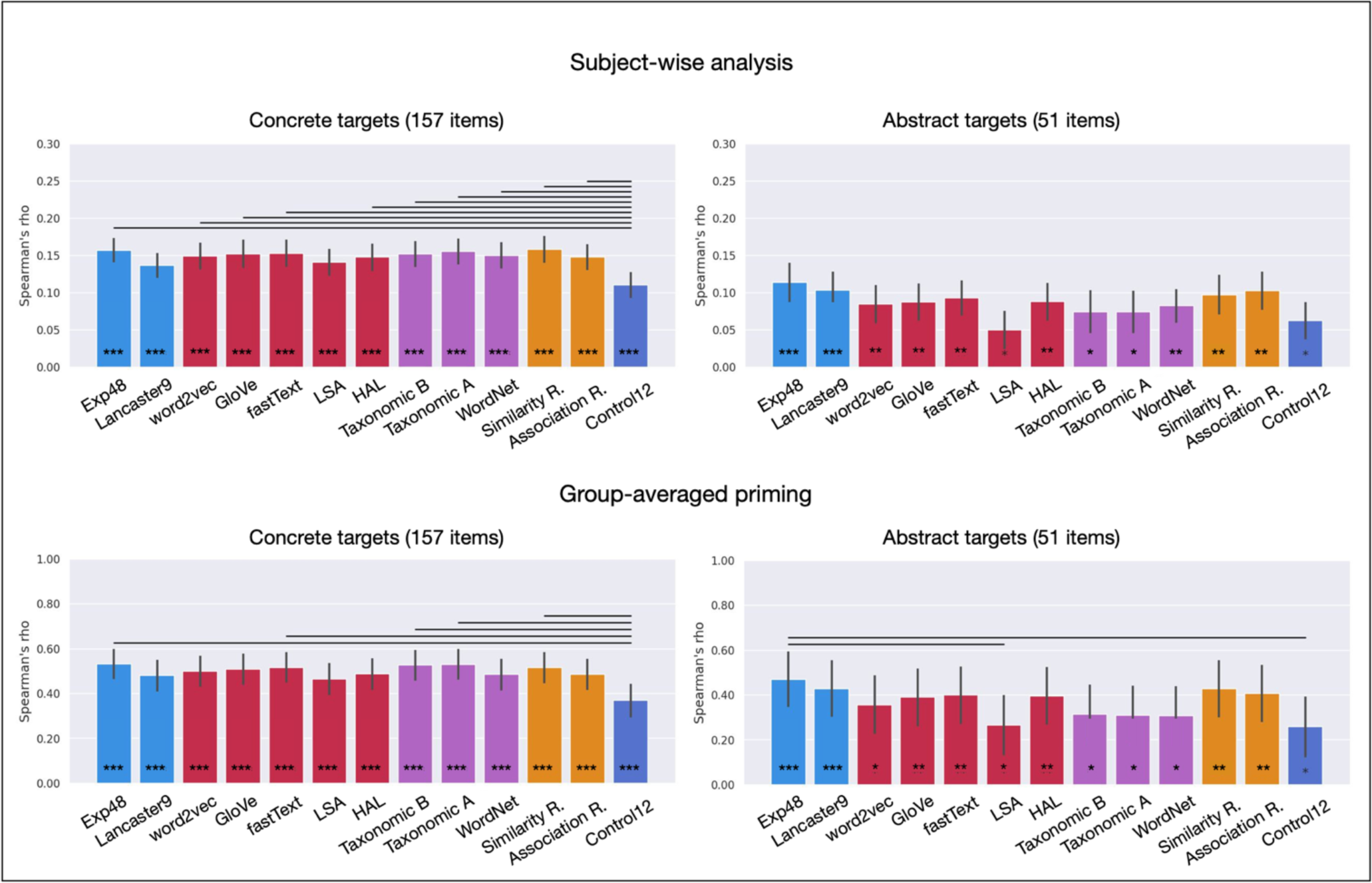
RSA correlations for experiential (light blue), distributional (red), and taxonomic (purple) models of semantic representation, for explicit ratings of semantic similarity and thematic association (orange), and for Control12 (dark blue). ***FDR-corrected p < .0005; **FDR- corrected p < .005; *FDR-corrected p < .05.

**Supplemental Figure 4.**
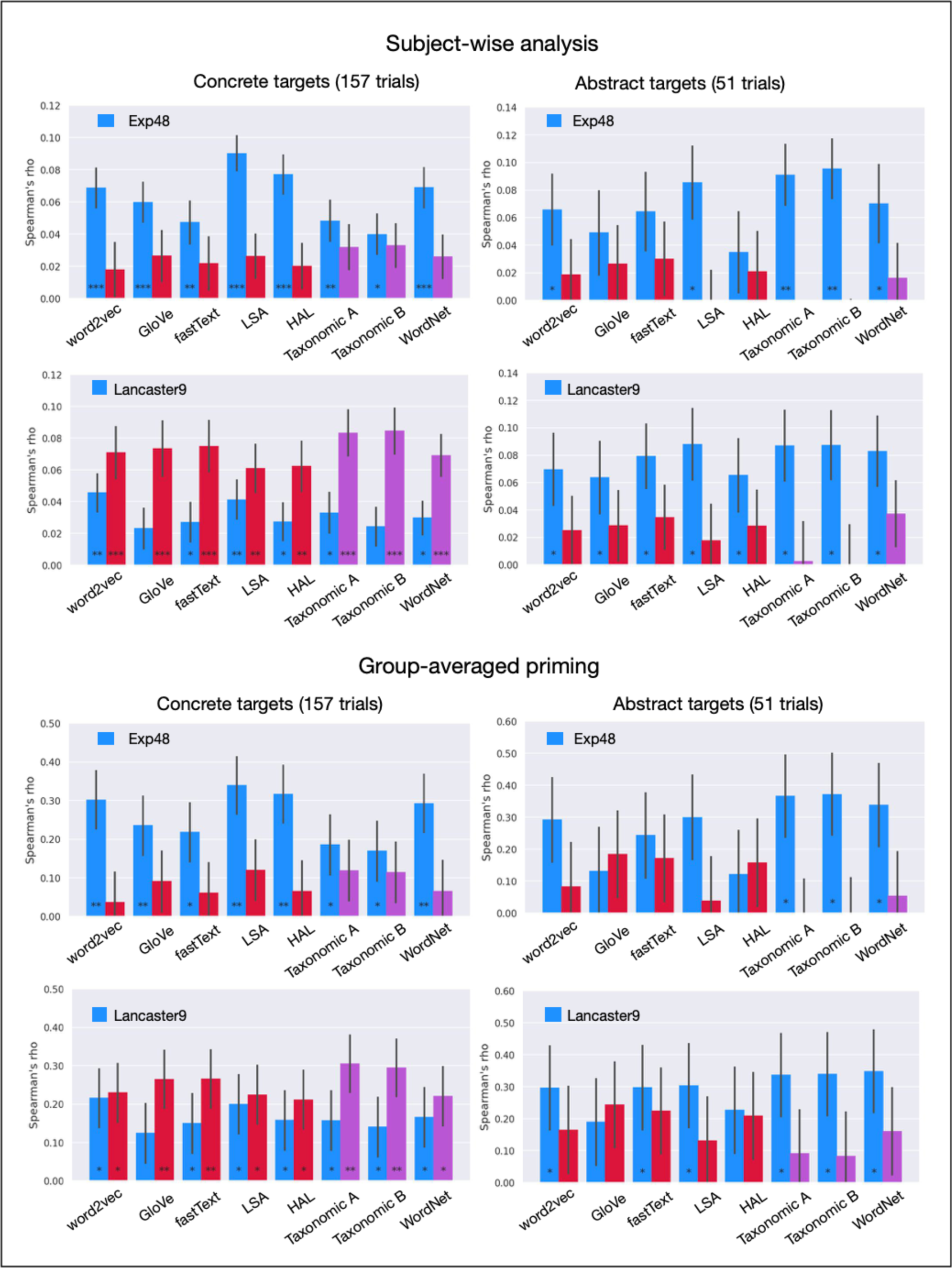
Pairwise partial correlations evaluating the unique variance explained by the experiential models (blue) while controlling for each of the other models, and the unique variance explained by the distributional (red) and taxonomic (purple) models while controlling for each experiential model. ***FDR-corrected p < .0005; **FDR-corrected p < .005; *FDR-corrected p < .05.

**Supplemental Figure 5.**
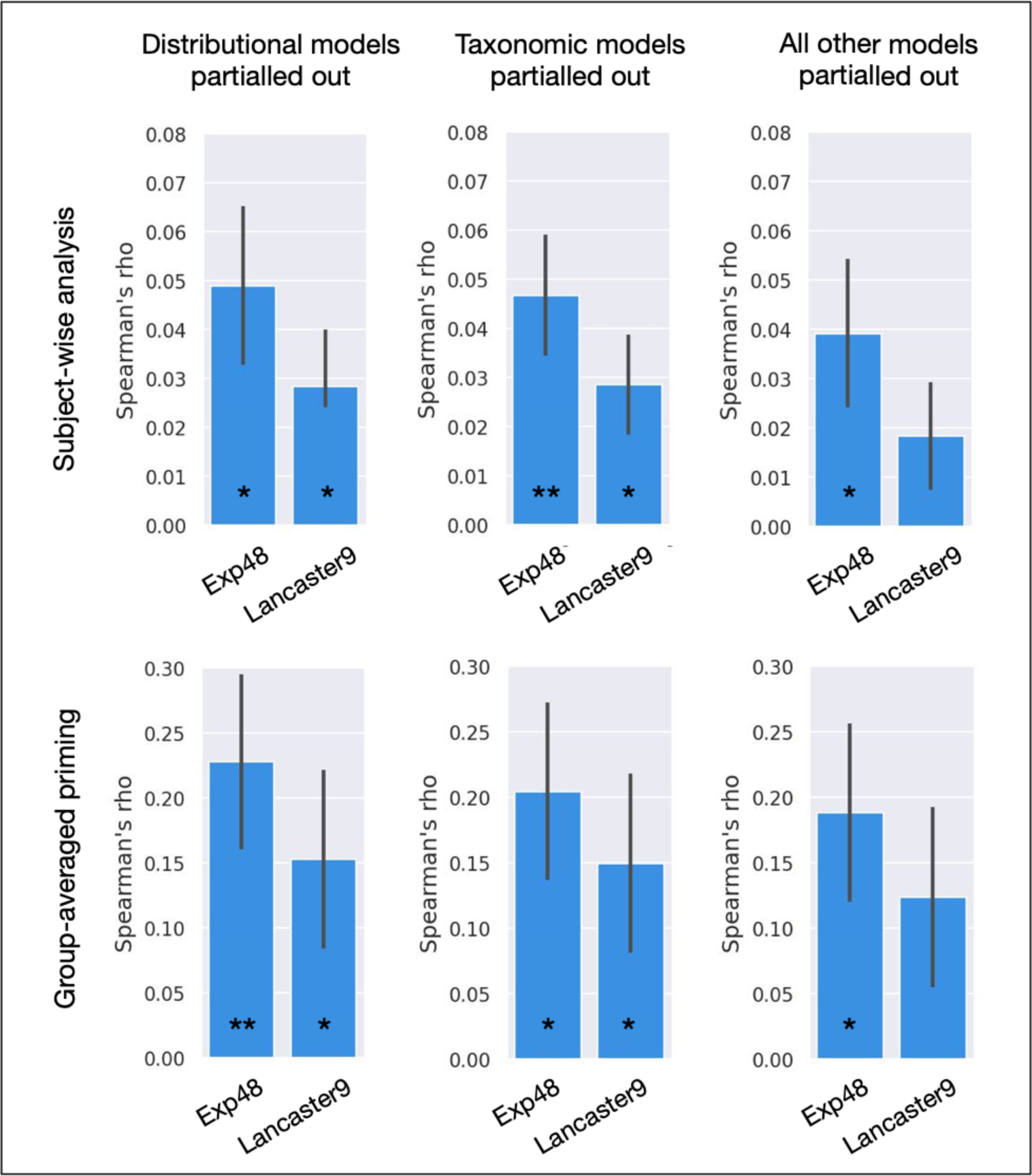
RSA partial correlations for each experiential model while partialling out the variance explained by all distributional models (left), all taxonomic models (center), and all distributional and taxonomic models simultaneously (right). **FDR-corrected p < .005; *FDR-corrected p < .05.

**Supplemental Figure 6.**
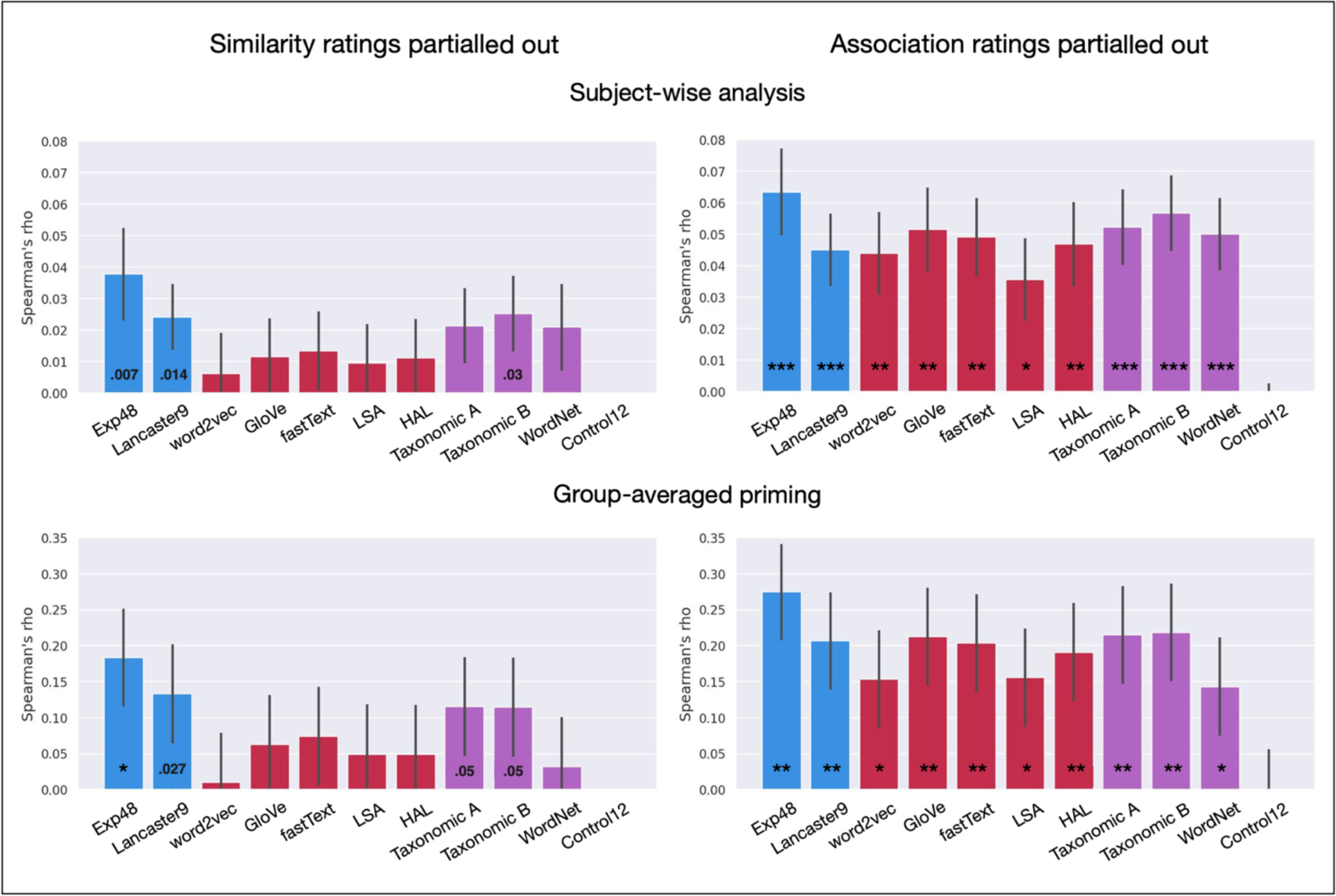
RSA correlations with the priming data after partialling out the similarity (left panels) or association (right panels) ratings. ***FDR-corrected p < .0005; **FDR-corrected p < .005; *FDR-corrected p < .05. Uncorrected p < .05 are shown as figures inside bars in the left-hand charts.

**Supplemental Table 1.** Word triplets used as stimuli in the study.

| Target | Session A Prime | Session B Prime |
| --- | --- | --- |
| accordion | whale | banjo |
| activist | teacher | election |
| alligator | snake | prison |
| ant | trust | mouse |
| apology | hand | truth |
| applause | rally | jury |
| apricot | animosity | tangerine |
| asparagus | cabbage | tornado |
| avalanche | chestnut | landslide |
| awe | sin | joy |
| axe | hoe | awe |
| bagpipe | saxophone | infinity |
| banana | luck | fish |
| banjo | parent | piano |
| banker | businessman | politician |
| barbecue | musical | dinner |
| bay | heroism | prairie |
| bed | crib | whine |
| bee | mosquito | clarinet |
| beer | tuba | rum |
| bell | drum | boat |
| boat | bus | beer |
| boy | child | night |
| bread | ball | ice |
| bribe | grievance | megaphone |
| broccoli | mushroom | butterfly |
| businessman | testimony | journalist |
| butterfly | turtle | penguin |
| cab | garden | plane |
| cabbage | plot | egg |
| cafeteria | store | summer |
| camel | jury | cab |
| car | clue | van |
| carnival | applause | toaster |
| carriage | ricochet | escalator |
| carrot | broccoli | gunshot |
| cathedral | company | office |
| cheetah | mercy | hawk |
| cherry | finger | coffee |
| chestnut | chocolate | television |
| chime | fiddle | pencil |
| chipmunk | screech | hamster |
| choir | audience | theater |
| circus | medicine | festival |
| comb | key | cash |
| corkscrew | hairbrush | cafeteria |
| corn | fireworks | pineapple |
| cough | belch | faucet |
| cranberry | cherry | deceit |
| crib | subway | shelves |
| crow | motive | monkey |
| cucumber | pumpkin | malice |
| cyclone | policeman | ambulance |
| dandelion | sailboat | raspberry |
| day | morning | evening |
| denial | treaty | farmer |
| dictation | oration | bonfire |
| dinner | criminal | kitchen |
| diplomat | editor | train |
| dog | spring | horse |
| dolphin | moose | muscle |
| downpour | moral | storm |
| dread | shame | bread |
| drum | highway | trumpet |
| duck | nose | pig |
| eggplant | cucumber | limousine |
| elm | council | mountain |
| embassy | school | college |
| embrace | quantity | vacation |
| envy | dread | cough |
| era | patent | mystery |
| etiquette | team | eye |
| farm | park | beach |
| farmer | pilot | excuse |
| fate | worth | power |
| fee | tax | tree |
| fiddle | accordion | stampede |
| fireworks | alligator | carnival |
| flute | camera | chicken |
| folly | fallacy | parade |
| foot | arm | fate |
| funeral | man | day |
| gasp | squeal | folly |
| goldfish | salmon | cloud |
| gratitude | peace | stone |
| grief | accident | agreement |
| grievance | rumor | denial |
| guilt | attribute | elevator |
| gum | cranberry | barbecue |
| gunshot | explosion | apartment |
| hailstorm | avalanche | intellect |
| hairbrush | umbrella | honeymoon |
| hall | driver | street |
| ham | spaghetti | delirium |
| hamster | bird | zone |
| harmonica | choir | flute |
| harp | xylophone | hailstorm |
| hawk | eggplant | cheetah |
| heredity | forest | reality |
| heroism | author | family |
| hoe | axe | gasp |
| honey | joke | jam |
| honeymoon | tourist | tribute |
| horse | dolphin | hygiene |
| hurricane | car | book |
| hygiene | knowledge | business |
| infinity | whole | woman |
| irony | shoulder | paradox |
| jam | ham | zoo |
| jaw | toe | lip |
| jealousy | ire | noun |
| joke | speech | student |
| journalist | activist | year |
| joy | gratitude | harmonica |
| jungle | airport | victim |
| jury | witness | bridge |
| ketchup | honey | rose |
| kiss | dread | satire |
| lab | embassy | cherry |
| lake | bay | fee |
| landslide | soccer | cyclone |
| law | legality | goldfish |
| leg | love | hand |
| legality | law | feather |
| limousine | jealousy | carriage |
| loan | door | semester |
| love | trust | finger |
| luck | advantage | hospital |
| lust | home | fun |
| malice | worker | torment |
| man | lab | girl |
| mandolin | harp | plea |
| mayor | commander | reporter |
| mob | role | army |
| monkey | bed | boy |
| moose | apology | elephant |
| mosquito | bee | sled |
| motive | couple | theory |
| mushroom | minister | newspaper |
| mustard | tomato | guilt |
| nose | hair | party |
| noun | analogy | embrace |
| oak | farm | elm |
| optimism | table | hope |
| pan | heredity | spatula |
| park | restaurant | terrorist |
| pen | truce | comb |
| pencil | pan | foot |
| penguin | scientist | minister |
| perjury | feather | excuse |
| pie | bicycle | cheese |
| pig | tiger | duck |
| pineapple | crow | corn |
| plea | window | debate |
| plot | trial | wonder |
| plum | dictation | blueberry |
| policeman | thunder | soldier |
| politician | year | water |
| prairie | lake | vice |
| pumpkin | carrot | volcano |
| radish | pea | rake |
| rake | delirium | corkscrew |
| rally | lawyer | protest |
| raspberry | gum | ant |
| riot | ticket | battle |
| river | symphony | fountain |
| role | tobacco | sympathy |
| rose | whine | tulip |
| rum | era | tea |
| salmon | apricot | ivy |
| sandpaper | downpour | stapler |
| saxophone | gong | bugle |
| scissors | loan | pen |
| scooter | lust | truck |
| screech | chair | scream |
| semester | football | baseball |
| shame | guilt | desk |
| shelves | desk | hall |
| sin | deceit | banker |
| snake | sun | dog |
| spaghetti | pie | bribe |
| spatula | patient | cabinet |
| squeal | scream | chair |
| stampede | hurricane | submarine |
| stapler | funeral | scissors |
| store | theme | hotel |
| sum | problem | number |
| tangerine | banana | jungle |
| tax | mercy | verb |
| tea | lemonade | ketchup |
| tiger | lightning | keyboard |
| toe | fee | leg |
| torment | riot | rally |
| tornado | glass | flood |
| tourist | belief | guard |
| tribute | meeting | engineer |
| trombone | bell | judge |
| truck | river | island |
| tuba | trombone | sandpaper |
| tulip | dandelion | chipmunk |
| turtle | sum | oak |
| van | optimism | scooter |
| verb | winter | advice |
| vice | envy | curse |
| voter | irony | mayor |
| whale | camel | voter |
| worker | mouth | priest |
| xylophone | mandolin | diplomat |
| zone | court | church |
| zoo | circus | flower |

**Supplemental Table 2.** Pooled lexical data for primes in each testing session. Prime sets were matched on all 24 lexical variables.

|  | Session A |  | Session B |  |
| --- | --- | --- | --- | --- |
|  | Mean | SE | Mean | SE |
| Freq_HAL | 8.80 | 0.12 | 8.78 | 0.12 |
| Freq_SUBTLEXUS | 2.82 | 0.05 | 2.88 | 0.05 |
| CD_SUBTLEXUS | 2.57 | 0.05 | 2.61 | 0.05 |
| CD_News | 2.732 | 0.06 | 2.726 | 0.06 |
| Fam_Brys | 0.997 | 0.00 | 0.997 | 0.00 |
| Prevalence_Brys | 2.31 | 0.02 | 2.34 | 0.01 |
| AoA_Kuper | 7 | 0.18 | 7 | 0.17 |
| NLett | 6 | 0.13 | 6 | 0.13 |
| BigramF_Avg_C | 1070 | 71.2 | 1045 | 91.1 |
| TrigramF_Avg_C | 239 | 25.8 | 217 | 21.6 |
| Levenshtein distance | 7 | 0.13 | 7 | 0.12 |
| Orth_N | 4.90 | 0.48 | 4.57 | 0.41 |
| NPhon | 4.96 | 0.12 | 4.96 | 0.13 |
| NSyll | 1.91 | 0.06 | 1.88 | 0.06 |
| UniphonP_Un_C | 0.065 | 0.001 | 0.063 | 0.00 |
| UniphonP_St_C | 0.061 | 0.001 | 0.059 | 0.00 |
| Phon_N | 11.4 | 1.09 | 9.9 | 0.90 |
| Nsenses_WordNet | 4.56 | 0.25 | 4.49 | 0.25 |
| Nmeanings_Websters | 3.40 | 0.17 | 3.24 | 0.17 |
| Sem_Diversity | 1.60 | 0.02 | 1.59 | 0.02 |
| NMorph | 1.30 | 0.04 | 1.30 | 0.04 |
| LexicalID_RT_V_ELP | 652 | 5.53 | 642 | 5.10 |
| Naming_RT_ELP | 633 | 3.87 | 628 | 3.34 |
| Recog_Memory | 0.54 | 0.01 | 0.56 | 0.01 |

**Supplemental Table 3.** Dimensions included in Exp48.

| Dimension | Rating query<br>(to what degree do you think of this thing as...) | High Score Examples |
| --- | --- | --- |
| Vision | being something you can easily see | moon, locomotive |
| Color | having a characteristic or defining color | grass, banana |
| Pattern | having a characteristic visual texture or pattern | tiger, pineapple |
| Bright | being visually light or bright | sun, lightning |
| Dark | being visually dark | night, crow |
| Motion | having a lot of visually observable movement | tornado, parade |
| Fast | having visible movement that is fast | rocket, cheetah |
| Slow | having visible movement that is slow | snail, cloud |
| Shape | having a characteristic visual shape or form | giraffe, spoon |
| Large | being large in size | volcano, ferry |
| Small | being small in size | bacterium, pea |
| Touch | easily recognizable by touch | toothbrush, sandpaper |
| Temperature | being either hot or cold to the touch | bonfire, ice |
| Texture | having either a smooth or rough texture to the touch | silk, stubble |
| Weight | being either light or heavy in weight | balloon, anvil |
| Pain | being associated with pain or physical discomfort | headache, bombing |
| Taste | having a characteristic taste | lemon, chocolate |
| Smell | having a characteristic smell | tobacco, barbecue |
| Mouth Action | associated with actions of the mouth | whistle, jaw |
| Hand Action | associated with actions of the arm, hand or fingers | keyboard, scissors |
| Foot Action | associated with actions of the legs or feet | soccer, bicycle |
| Manipulation | a physical object you have personally manipulated | fork, computer |
| Landmark | having a fixed location, as on a map | airport, library |
| Near | being near you, within reaching distance | foot, chair |
| Scene | belonging to a particular setting or physical location | oven, beach |
| Sound | having a characteristic or recognizable sound | rooster, piano |
| Audition | being something that you can easily hear | siren, thunder |
| Loud | making a loud sound | explosion, megaphone |
| Low pitch | having a low-pitched sound | tuba, growling |
| High pitch | having a high-pitched sound | whistle, dolphin |
| Trajectory | showing motion along a particular direction or path | rocket, trolley |
| Approaching | being associated with movement toward you | food, embrace |
| Receding | being associated with movement away from you | cough, kite |
| Time | an event that occurs at a typical or predictable time | lunch, night |
| Duration | having a predictable duration, whether short or long | movie, year |
| Long-lasting | an event that lasts for a long period of time | life, infinity |
| Short-lasting | an event that lasts for a short period of time | sneeze, gunshot |
| Caused | caused by a preceding event, action, or situation | spill, honeymoon |
| Consequential | likely to have consequences | invasion, earthquake |
| Intention | having human-like intentions, plans, or goals | lobbyist, activist |
| Benefit | something that could help or benefit you or others | cure, peace |
| Harm | something that could cause harm to you or others | epidemic, wildfire |
| Drive | something that motivates you to do something | duty, hope |
| Needs | something that would be hard to live without | shelter, water |
| Attention | something that grabs your attention | scream, lightning |
| Pleasant | something that you find pleasant | vacation, cake |
| Unpleasant | something that you find unpleasant | pain, bombing |
| Arousal | something that makes you feel alert, excited, or keyed-up | rollercoaster, lust |

**Supplemental Table 4.**
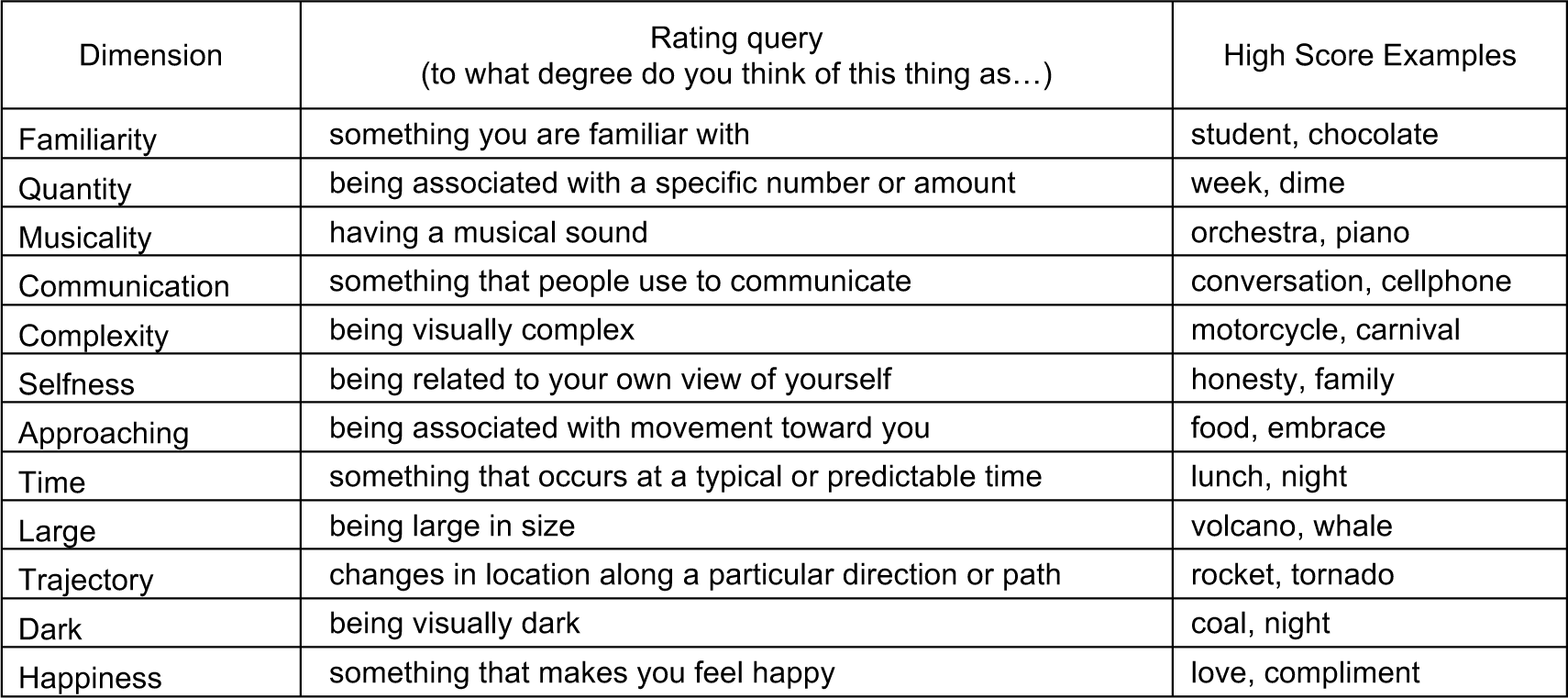
Dimensions included in the Control12 model.

**Supplemental Table 5.** Pairwise correlations between priming predictions from all semantic models and ratings in the main analysis (including all trials).

|  | Exp48 | Lancaster | word2vec | GloVe | fastText | LSA | HAL | TaxonA | TaxonB | WordNet | SimilarityR. |
| --- | --- | --- | --- | --- | --- | --- | --- | --- | --- | --- | --- |
| Lancaster | 0.80 |  |  |  |  |  |  |  |  |  |  |
| word2vec | 0.82 | 0.68 |  |  |  |  |  |  |  |  |  |
| GloVe | 0.85 | 0.74 | 0.93 |  |  |  |  |  |  |  |  |
| fastText | 0.86 | 0.74 | 0.94 | 0.94 |  |  |  |  |  |  |  |
| LSA | 0.73 | 0.66 | 0.83 | 0.91 | 0.81 |  |  |  |  |  |  |
| HAL | 0.83 | 0.75 | 0.89 | 0.95 | 0.89 | 0.94 |  |  |  |  |  |
| CategoricalA | 0.86 | 0.68 | 0.79 | 0.80 | 0.82 | 0.70 | 0.77 |  |  |  |  |
| CategoricalB | 0.88 | 0.70 | 0.82 | 0.82 | 0.84 | 0.72 | 0.80 | 0.99 |  |  |  |
| WordNet | 0.77 | 0.67 | 0.78 | 0.78 | 0.78 | 0.67 | 0.76 | 0.77 | 0.80 |  |  |
| Similarity ratings | 0.91 | 0.76 | 0.90 | 0.92 | 0.93 | 0.80 | 0.88 | 0.87 | 0.89 | 0.85 |  |
| Association ratings | 0.84 | 0.69 | 0.81 | 0.86 | 0.85 | 0.75 | 0.82 | 0.77 | 0.78 | 0.74 | 0.90 |

**Supplemental Table 6.** Pairwise correlations between priming predictions from all semantic models and ratings in the subset of trials for which SFPN norms are available.

|  | Exp48 | Lancaster | word2vec | GloVe | fastText | LSA | HAL | TaxonA | TaxonB | WordNet | Simil.R. | Assoc.R. |
| --- | --- | --- | --- | --- | --- | --- | --- | --- | --- | --- | --- | --- |
| Lancaster | 0.79 |  |  |  |  |  |  |  |  |  |  |  |
| word2vec | 0.83 | 0.64 |  |  |  |  |  |  |  |  |  |  |
| GloVe | 0.84 | 0.72 | 0.93 |  |  |  |  |  |  |  |  |  |
| fastText | 0.88 | 0.71 | 0.95 | 0.94 |  |  |  |  |  |  |  |  |
| LSA | 0.70 | 0.63 | 0.83 | 0.92 | 0.81 |  |  |  |  |  |  |  |
| HAL | 0.78 | 0.72 | 0.88 | 0.96 | 0.88 | 0.95 |  |  |  |  |  |  |
| CategoricalA | 0.81 | 0.66 | 0.76 | 0.75 | 0.78 | 0.63 | 0.70 |  |  |  |  |  |
| CategoricalB | 0.84 | 0.69 | 0.81 | 0.79 | 0.82 | 0.66 | 0.75 | 0.98 |  |  |  |  |
| WordNet | 0.76 | 0.65 | 0.74 | 0.75 | 0.76 | 0.63 | 0.73 | 0.75 | 0.80 |  |  |  |
| Similarity ratings | 0.90 | 0.75 | 0.90 | 0.90 | 0.92 | 0.79 | 0.87 | 0.84 | 0.89 | 0.85 |  |  |
| Association ratings | 0.85 | 0.68 | 0.80 | 0.83 | 0.84 | 0.71 | 0.78 | 0.73 | 0.76 | 0.77 | 0.90 |  |
| SFPN (McRae features) | 0.61 | 0.57 | 0.73 | 0.74 | 0.73 | 0.71 | 0.73 | 0.55 | 0.60 | 0.67 | 0.74 | 0.66 |

**Supplemental Table 7.** Results of ordinary least squares models of group-averaged priming including one regressor (predictions from a single semantic model), analogous to the main RSA (sorted by increasing AIC).

| Regressor | AIC | BIC | LogLikelihood | p | R<sup>2</sup> |
| --- | --- | --- | --- | --- | --- |
| Exp48: | 22.1 | 28.8 | -9.04 | **8.71E-16** | 0.27 |
| SimilarityR: | 27.8 | 34.5 | -11.92 | **1.57E-14** | 0.25 |
| GloVe: | 32.8 | 39.4 | -14.38 | **1.85E-13** | 0.23 |
| FastText: | 33.1 | 39.8 | -14.54 | **2.18E-13** | 0.23 |
| TaxonomicB: | 34.3 | 41 | -15.17 | **4.12E-13** | 0.22 |
| TaxonomicA: | 34.5 | 41.2 | -15.26 | **4.53E-13** | 0.22 |
| Lancaster: | 37.2 | 43.9 | -16.58 | **1.71E-12** | 0.21 |
| HAL: | 37.3 | 44 | -16.64 | **1.82E-12** | 0.21 |
| AssociationR: | 37.5 | 44.2 | -16.77 | **2.07E-12** | 0.21 |
| word2vec: | 39.2 | 45.9 | -17.59 | **4.71E-12** | 0.21 |
| WordNet: | 41.3 | 48 | -18.67 | **1.40E-11** | 0.20 |
| LSA: | 46.4 | 53.1 | -21.2 | **1.80E-10** | 0.18 |
| Control12: | 64.2 | 70.8 | -30.08 | **1.52E-06** | 0.11 |

**Supplemental Table 8.** Results of the linear model AIC comparisons for pairs of semantic model predictions. For each semantic model and for each participant, a linear regression model was fit using the model’s priming predictions as the predictor variable, producing an AIC index. The AIC difference between two models was tested for significance across subjects with Wilcoxon’s signed rank tests (two-tailed, n = 31).

| AIC comparison (signed rank test) | p |
| --- | --- |
| Exp48 vs. Lancaster9: | 0.053 |
| Exp48 vs. word2vec: | 0.195 |
| Exp48 vs. GloVe: | 0.308 |
| Exp48 vs. FastText: | 0.318 |
| Exp48 vs. LSA: | **0.018** |
| Exp48 vs. HAL: | 0.195 |
| Exp48 vs. TaxonomicA: | 0.318 |
| Exp48 vs. TaxonomicB: | 0.337 |
| Exp48 vs. WordNet: | 0.058 |
| Exp48 vs. SimilarityR: | 0.357 |
| Exp48 vs. AssociationR: | **0.036** |
| Exp48 vs. Control12: | **0.000** |
| Lancaster9 vs. Exp48: | 0.053 |
| Lancaster9 vs. word2vec: | 0.854 |
| Lancaster9 vs. GloVe: | 0.410 |
| Lancaster9 vs. FastText: | 0.468 |
| Lancaster9 vs. LSA: | 0.608 |
| Lancaster9 vs. HAL: | 0.480 |
| Lancaster9 vs. TaxonomicA: | 1 |
| Lancaster9 vs. TaxonomicB: | 0.900 |
| Lancaster9 vs. WordNet: | 0.931 |
| Lancaster9 vs. SimilarityR: | 0.468 |
| Lancaster9 vs. AssociationR: | 0.870 |
| Lancaster9 vs. Control12: | **0.020** |
| word2vec vs. Exp48: | 0.195 |
| word2vec vs. Lancaster9: | 0.854 |
| word2vec vs. GloVe: | 0.102 |
| word2vec vs. FastText: | 0.169 |
| word2vec vs. LSA: | 0.378 |
| word2vec vs. HAL: | 0.327 |
| word2vec vs. TaxonomicA: | 0.839 |
| word2vec vs. TaxonomicB: | 0.664 |
| word2vec vs. WordNet: | 0.765 |
| word2vec vs. SimilarityR: | 0.256 |
| word2vec vs. AssociationR: | 0.542 |
| word2vec vs. Control12: | **0.021** |
| GloVe vs. Exp48: | 0.308 |
| GloVe vs. Lancaster9: | 0.41 |
| GloVe vs. word2vec: | 0.102 |
| GloVe vs. FastText: | 0.946 |
| GloVe vs. LSA: | **0.004** |
| GloVe vs. HAL: | 0.224 |
| GloVe vs. TaxonomicA: | 0.232 |
| GloVe vs. TaxonomicB: | 0.256 |
| GloVe vs. WordNet: | 0.195 |
| GloVe vs. SimilarityR: | 0.961 |
| GloVe vs. AssociationR: | 0.195 |
| GloVe vs. Control12: | **0.004** |
| FastText vs. Exp48: | 0.318 |
| FastText vs. Lancaster9: | 0.468 |
| FastText vs. word2vec: | 0.169 |
| FastText vs. GloVe: | 0.946 |
| FastText vs. LSA: | **0.017** |
| FastText vs. HAL: | 0.125 |
| FastText vs. TaxonomicA: | 0.399 |
| FastText vs. TaxonomicB: | 0.399 |
| FastText vs. WordNet: | 0.367 |
| FastText vs. SimilarityR: | 0.75 |
| FastText vs. AssociationR: | 0.157 |
| FastText vs. Control12: | **0.002** |
| LSA vs. Exp48: | **0.018** |
| LSA vs. Lancaster9: | 0.608 |
| LSA vs. word2vec: | 0.378 |
| LSA vs. GloVe: | **0.004** |
| LSA vs. FastText: | **0.017** |
| LSA vs. HAL: | 0.098 |
| LSA vs. TaxonomicA: | 0.517 |
| LSA vs. TaxonomicB: | 0.264 |
| LSA vs. WordNet: | 0.327 |
| LSA vs. SimilarityR: | **0.030** |
| LSA vs. AssociationR: | 0.492 |
| LSA vs. Control12: | **0.021** |
| HAL vs. Exp48: | 0.195 |
| HAL vs. Lancaster9: | 0.480 |
| HAL vs. word2vec: | 0.327 |
| HAL vs. GloVe: | 0.224 |
| HAL vs. FastText: | 0.125 |
| HAL vs. LSA: | 0.098 |
| HAL vs. TaxonomicA: | 0.529 |
| HAL vs. TaxonomicB: | 0.595 |
| HAL vs. WordNet: | 0.542 |
| HAL vs. SimilarityR: | 0.542 |
| HAL vs. AssociationR: | 0.706 |
| HAL vs. Control12: | **0.005** |
| TaxonomicA vs. Exp48: | 0.318 |
| TaxonomicA vs. Lancaster9: | 1 |
| TaxonomicA vs. word2vec: | 0.839 |
| TaxonomicA vs. GloVe: | 0.232 |
| TaxonomicA vs. FastText: | 0.399 |
| TaxonomicA vs. LSA: | 0.517 |
| TaxonomicA vs. HAL: | 0.529 |
| TaxonomicA vs. TaxonomicB: | 0.650 |
| TaxonomicA vs. WordNet: | 0.870 |
| TaxonomicA vs. SimilarityR: | 0.299 |
| TaxonomicA vs. AssociationR: | 0.444 |
| TaxonomicA vs. Control12: | **0.010** |
| TaxonomicB vs. Exp48: | 0.337 |
| TaxonomicB vs. Lancaster9: | 0.900 |
| TaxonomicB vs. word2vec: | 0.664 |
| TaxonomicB vs. GloVe: | 0.256 |
| TaxonomicB vs. FastText: | 0.399 |
| TaxonomicB vs. LSA: | 0.264 |
| TaxonomicB vs. HAL: | 0.595 |
| TaxonomicB vs. TaxonomicA: | 0.650 |
| TaxonomicB vs. WordNet: | 0.721 |
| TaxonomicB vs. SimilarityR: | 0.347 |
| TaxonomicB vs. AssociationR: | 0.318 |
| TaxonomicB vs. Control12: | **0.004** |
| WordNet vs. Exp48: | 0.058 |
| WordNet vs. Lancaster9: | 0.931 |
| WordNet vs. word2vec: | 0.765 |
| WordNet vs. GloVe: | 0.195 |
| WordNet vs. FastText: | 0.367 |
| WordNet vs. LSA: | 0.327 |
| WordNet vs. HAL: | 0.542 |
| WordNet vs. TaxonomicA: | 0.870 |
| WordNet vs. TaxonomicB: | 0.721 |
| WordNet vs. SimilarityR: | 0.146 |
| WordNet vs. AssociationR: | 0.915 |
| WordNet vs. Control12: | 0.094 |
| SimilarityR vs. Exp48: | 0.357 |
| SimilarityR vs. Lancaster9: | 0.468 |
| SimilarityR vs. word2vec: | 0.256 |
| SimilarityR vs. GloVe: | 0.961 |
| SimilarityR vs. FastText: | 0.750 |
| SimilarityR vs. LSA: | **0.030** |
| SimilarityR vs. HAL: | 0.542 |
| SimilarityR vs. TaxonomicA: | 0.299 |
| SimilarityR vs. TaxonomicB: | 0.347 |
| SimilarityR vs. WordNet: | 0.146 |
| SimilarityR vs. AssociationR: | **0.014** |
| SimilarityR vs. Control12: | **0.000** |
| AssociationR vs. Exp48: | **0.036** |
| AssociationR vs. Lancaster9: | 0.870 |
| AssociationR vs. word2vec: | 0.542 |
| AssociationR vs. GloVe: | 0.195 |
| AssociationR vs. FastText: | 0.157 |
| AssociationR vs. LSA: | 0.492 |
| AssociationR vs. HAL: | 0.706 |
| AssociationR vs. TaxonomicA: | 0.444 |
| AssociationR vs. TaxonomicB: | 0.318 |
| AssociationR vs. WordNet: | 0.915 |
| AssociationR vs. SimilarityR: | **0.014** |
| AssociationR vs. Control12: | **0.013** |
| Control12 vs. Exp48: | **0.000** |
| Control12 vs. Lancaster9: | **0.020** |
| Control12 vs. word2vec: | **0.021** |
| Control12 vs. GloVe: | **0.004** |
| Control12 vs. FastText: | **0.002** |
| Control12 vs. LSA: | **0.021** |
| Control12 vs. HAL: | **0.005** |
| Control12 vs. TaxonomicA: | **0.010** |
| Control12 vs. TaxonomicB: | **0.004** |
| Control12 vs. WordNet: | 0.094 |
| Control12 vs. SimilarityR: | **0.000** |
| Control12 vs. AssociationR: | **0.013** |

**Supplemental Table 9.** Results of ordinary least squares models including two regressors (predictions from two semantic models), analogous to the pairwise partial correlation analyses.

|  | Target model<br>p | Control model<br>p | R<sup>2</sup> | AIC | BIC | LogLikelihood |
| --- | --- | --- | --- | --- | --- | --- |
| Exp48 controlled for: |  |  |  |  |  |  |
| Lancaster9 | **4.85E-05** | 1.97E-01 | 0.27 | 22.38 | 32.42 | -8.19 |
| word2vec | **2.95E-05** | 4.26E-01 | 0.27 | 23.43 | 33.47 | -8.72 |
| GloVe | **5.42E-04** | 2.26E-01 | 0.27 | 22.58 | 32.63 | -8.29 |
| FastText | **5.68E-04** | 3.01E-01 | 0.27 | 22.99 | 33.03 | -8.5 |
| LSA | **5.77E-07** | 2.99E-01 | 0.27 | 22.98 | 33.02 | -8.49 |
| HAL | **6.78E-05** | 3.39E-01 | 0.27 | 23.15 | 33.19 | -8.57 |
| TaxonomicA | **2.94E-04** | 3.53E-01 | 0.27 | 23.2 | 33.24 | -8.6 |
| TaxonomicB | **4.15E-04** | 5.26E-01 | 0.27 | 23.67 | 33.71 | -8.83 |
| WordNet | **6.63E-06** | 2.45E-01 | 0.27 | 22.7 | 32.74 | -8.35 |
| SimilarityR | **9.19E-03** | 2.91E-01 | 0.27 | 22.94 | 32.99 | -8.47 |
| AssociationR | **6.93E-05** | 4.32E-01 | 0.27 | 23.45 | 33.49 | -8.73 |
| Control12 | **9.30E-14** | **1.71E-04** | 0.32 | 9.71 | 19.75 | -1.85 |
| Lancaster9 controlled for: |  |  |  |  |  |  |
| Exp48 | 1.97E-01 | **4.85E-05** | 0.27 | 22.38 | 32.42 | -8.19 |
| word2vec | **6.31E-04** | **1.83E-03** | 0.25 | 29.3 | 39.34 | -11.65 |
| GloVe | **9.36E-03** | **8.64E-04** | 0.25 | 27.88 | 37.93 | -10.94 |
| FastText | **7.23E-03** | **7.98E-04** | 0.26 | 27.73 | 37.78 | -10.87 |
| LSA | **8.47E-05** | **1.15E-02** | 0.24 | 32.68 | 42.72 | -13.34 |
| HAL | **3.83E-03** | **4.08E-03** | 0.24 | 30.78 | 40.82 | -12.39 |
| TaxonomicA | **1.50E-03** | **3.71E-04** | 0.26 | 26.28 | 36.32 | -10.14 |
| TaxonomicB | **2.56E-03** | **5.67E-04** | 0.26 | 27.09 | 37.13 | -10.54 |
| WordNet | **3.49E-04** | **3.22E-03** | 0.25 | 30.34 | 40.38 | -12.17 |
| SimilarityR | **3.47E-02** | **2.22E-04** | 0.26 | 25.31 | 35.35 | -9.65 |
| AssociationR | **1.21E-03** | **1.48E-03** | 0.25 | 28.9 | 38.94 | -11.45 |
| Control12 | **1.67E-07** | 3.57E-01 | 0.22 | 38.31 | 48.35 | -16.15 |
| word2vec controlled for: |  |  |  |  |  |  |
| Exp48 | 4.26E-01 | **2.95E-05** | 0.27 | 23.43 | 33.47 | -8.72 |
| Lancaster9 | **1.83E-03** | **6.31E-04** | 0.25 | 29.3 | 39.34 | -11.65 |
| GloVe | 7.26E-01 | **1.12E-02** | 0.23 | 34.63 | 44.67 | -14.31 |
| FastText | 8.69E-01 | **1.40E-02** | 0.23 | 35.04 | 45.08 | -14.52 |
| LSA | **2.83E-03** | **1.78E-01** | 0.21 | 39.33 | 49.37 | -16.66 |
| HAL | 1.19E-01 | **3.83E-02** | 0.22 | 36.81 | 46.85 | -15.41 |
| TaxonomicA | **3.04E-02** | **2.35E-03** | 0.24 | 31.77 | 41.81 | -12.88 |
| TaxonomicB | 5.66E-02 | **3.77E-03** | 0.24 | 32.64 | 42.68 | -13.32 |
| WordNet | **6.17E-03** | **2.05E-02** | 0.23 | 35.72 | 45.76 | -14.86 |
| SimilarityR | 8.37E-01 | **8.24E-04** | 0.25 | 29.8 | 39.84 | -11.9 |
| AssociationR | **2.65E-02** | **1.06E-02** | 0.23 | 34.54 | 44.58 | -14.27 |
| Control12 | **7.17E-07** | 9.38E-01 | 0.21 | 41.17 | 51.21 | -17.59 |
| GloVe controlled for: |  |  |  |  |  |  |
| Exp48 | 2.26E-01 | **5.42E-04** | 0.27 | 22.58 | 32.63 | -8.29 |
| Lancaster9 | **8.64E-04** | **9.36E-03** | 0.25 | 27.88 | 37.93 | -10.94 |
| word2vec | **1.12E-02** | 7.26E-01 | 0.23 | 34.63 | 44.67 | -14.31 |
| FastText | 1.67E-01 | 2.07E-01 | 0.24 | 33.13 | 43.17 | -13.57 |
| LSA | **2.13E-04** | 5.86E-01 | 0.23 | 34.45 | 44.49 | -14.22 |
| HAL | **3.37E-02** | 8.25E-01 | 0.23 | 34.7 | 44.74 | -14.35 |
| TaxonomicA | **5.13E-03** | **1.36E-02** | 0.25 | 28.56 | 38.61 | -11.28 |
| TaxonomicB | **9.15E-03** | **2.22E-02** | 0.25 | 29.43 | 39.47 | -11.72 |
| WordNet | **5.88E-04** | 6.66E-02 | 0.24 | 31.33 | 41.37 | -12.66 |
| SimilarityR | 3.20E-01 | **1.59E-02** | 0.25 | 28.84 | 38.88 | -11.42 |
| AssociationR | **7.61E-03** | 1.21E-01 | 0.24 | 32.3 | 42.35 | -13.15 |
| Control12 | **2.44E-08** | 6.34E-01 | 0.23 | 34.52 | 44.56 | -14.26 |
| FastText controlled for: |  |  |  |  |  |  |
| Exp48 | 3.01E-01 | **5.68E-04** | 0.27 | 22.99 | 33.03 | -8.5 |
| Lancaster9 | **7.98E-04** | **7.23E-03** | 0.26 | 27.73 | 37.78 | -10.87 |
| word2vec | **1.40E-02** | 8.69E-01 | 0.23 | 35.04 | 45.08 | -14.52 |
| GloVe | 2.07E-01 | 1.67E-01 | 0.24 | 33.13 | 43.17 | -13.57 |
| LSA | **1.88E-04** | 3.58E-01 | 0.23 | 34.21 | 44.25 | -14.11 |
| HAL | **1.49E-02** | 1.82E-01 | 0.24 | 33.26 | 43.3 | -13.63 |
| TaxonomicA | **8.63E-03** | **1.94E-02** | 0.25 | 29.51 | 39.56 | -11.76 |
| TaxonomicB | **1.44E-02** | **2.94E-02** | 0.25 | 30.25 | 40.29 | -12.12 |
| WordNet | **6.08E-04** | 5.72E-02 | 0.24 | 31.39 | 41.43 | -12.7 |
| SimilarityR | 4.40E-01 | **1.66E-02** | 0.25 | 29.24 | 39.28 | -11.62 |
| AssociationR | **6.96E-03** | 8.96E-02 | 0.24 | 32.14 | 42.18 | -13.07 |
| Control12 | **2.68E-08** | 5.47E-01 | 0.23 | 34.7 | 44.74 | -14.35 |
| LSA controlled for: |  |  |  |  |  |  |
| Exp48 | 2.99E-01 | **5.77E-07** | 0.27 | 22.98 | 33.02 | -8.49 |
| Lancaster9 | **1.15E-02** | **8.47E-05** | 0.24 | 32.68 | 42.72 | -13.34 |
| word2vec | 1.78E-01 | **2.83E-03** | 0.21 | 39.33 | 49.37 | -16.66 |
| GloVe | 5.86E-01 | **2.13E-04** | 0.23 | 34.45 | 44.49 | -14.22 |
| FastText | 3.58E-01 | **1.88E-04** | 0.23 | 34.21 | 44.25 | -14.11 |
| HAL | 6.23E-01 | **2.43E-03** | 0.21 | 39.04 | 49.08 | -16.52 |
| TaxonomicA | **3.41E-02** | **5.84E-05** | 0.24 | 31.96 | 42.01 | -12.98 |
| TaxonomicB | 5.28E-02 | **7.82E-05** | 0.24 | 32.53 | 42.57 | -13.26 |
| WordNet | **6.58E-03** | **4.41E-04** | 0.23 | 35.83 | 45.87 | -14.92 |
| SimilarityR | 5.24E-01 | **1.58E-05** | 0.25 | 29.43 | 39.47 | -11.72 |
| AssociationR | 6.34E-02 | **4.93E-04** | 0.22 | 36.04 | 46.08 | -15.02 |
| Control12 | **1.18E-05** | 1.88E-01 | 0.18 | 46.63 | 56.67 | -20.31 |
| HAL controlled for: |  |  |  |  |  |  |
| Exp48 | 3.39E-01 | **6.78E-05** | 0.27 | 23.15 | 33.19 | -8.57 |
| Lancaster9 | **4.08E-03** | **3.83E-03** | 0.24 | 30.78 | 40.82 | -12.39 |
| word2vec | **3.83E-02** | 1.19E-01 | 0.22 | 36.81 | 46.85 | -15.41 |
| GloVe | 8.25E-01 | **3.37E-02** | 0.23 | 34.7 | 44.74 | -14.35 |
| FastText | 1.82E-01 | **1.49E-02** | 0.24 | 33.26 | 43.3 | -13.63 |
| LSA | **2.43E-03** | 6.23E-01 | 0.21 | 39.04 | 49.08 | -16.52 |
| TaxonomicA | **1.23E-02** | **2.72E-03** | 0.25 | 30.15 | 40.19 | -12.08 |
| TaxonomicB | **2.21E-02** | **4.33E-03** | 0.24 | 31.01 | 41.05 | -12.51 |
| WordNet | **1.90E-03** | **1.74E-02** | 0.23 | 33.54 | 43.58 | -13.77 |
| SimilarityR | 4.20E-01 | **1.62E-03** | 0.25 | 29.18 | 39.23 | -11.59 |
| AssociationR | **1.64E-02** | **1.90E-02** | 0.23 | 33.69 | 43.73 | -13.84 |
| Control12 | **2.74E-07** | 9.58E-01 | 0.21 | 39.28 | 49.33 | -16.64 |
| TaxonomicA controlled for: |  |  |  |  |  |  |
| Exp48 | 3.53E-01 | **2.94E-04** | 0.27 | 23.2 | 33.24 | -8.6 |
| Lancaster9 | **3.71E-04** | **1.50E-03** | 0.26 | 26.28 | 36.32 | -10.14 |
| word2vec | **2.35E-03** | **3.04E-02** | 0.24 | 31.77 | 41.81 | -12.88 |
| GloVe | **1.36E-02** | **5.13E-03** | 0.25 | 28.56 | 38.61 | -11.28 |
| FastText | **1.94E-02** | **8.63E-03** | 0.25 | 29.51 | 39.56 | -11.76 |
| LSA | **5.84E-05** | **3.41E-02** | 0.24 | 31.96 | 42.01 | -12.98 |
| HAL | **2.72E-03** | **1.23E-02** | 0.25 | 30.15 | 40.19 | -12.08 |
| TaxonomicB | 5.51E-01 | 4.62E-01 | 0.23 | 35.98 | 46.02 | -14.99 |
| WordNet | **9.21E-04** | **3.86E-02** | 0.24 | 32.18 | 42.22 | -13.09 |
| SimilarityR | 1.66E-01 | **3.56E-03** | 0.25 | 27.89 | 37.93 | -10.95 |
| AssociationR | **2.24E-03** | **1.16E-02** | 0.25 | 30.05 | 40.09 | -12.02 |
| Control12 | **6.24E-08** | 6.93E-01 | 0.22 | 36.37 | 46.41 | -15.19 |
| TaxonomicB controlled for: |  |  |  |  |  |  |
| Exp48 | 5.26E-01 | **4.15E-04** | 0.27 | 23.67 | 33.71 | -8.83 |
| Lancaster9 | **5.67E-04** | **2.56E-03** | 0.26 | 27.09 | 37.13 | -10.54 |
| word2vec | **3.77E-03** | 5.66E-02 | 0.24 | 32.64 | 42.68 | -13.32 |
| GloVe | **2.22E-02** | **9.15E-03** | 0.25 | 29.43 | 39.47 | -11.72 |
| FastText | **2.94E-02** | **1.44E-02** | 0.25 | 30.25 | 40.29 | -12.12 |
| LSA | **7.82E-05** | 5.28E-02 | 0.24 | 32.53 | 42.57 | -13.26 |
| HAL | **4.33E-03** | **2.21E-02** | 0.24 | 31.01 | 41.05 | -12.51 |
| TaxonomicA | 4.62E-01 | 5.51E-01 | 0.23 | 35.98 | 46.02 | -14.99 |
| WordNet | **1.51E-03** | 7.43E-02 | 0.24 | 33.1 | 43.14 | -13.55 |
| SimilarityR | 2.93E-01 | **6.20E-03** | 0.25 | 28.72 | 38.76 | -11.36 |
| AssociationR | **3.02E-03** | **1.75E-02** | 0.24 | 30.6 | 40.64 | -12.3 |
| Control12 | **4.47E-08** | 4.33E-01 | 0.23 | 35.71 | 45.76 | -14.86 |
| WordNet controlled for: |  |  |  |  |  |  |
| Exp48 | 2.45E-01 | **6.63E-06** | 0.27 | 22.7 | 32.74 | -8.35 |
| Lancaster9 | **3.22E-03** | **3.49E-04** | 0.25 | 30.34 | 40.38 | -12.17 |
| word2vec | **2.05E-02** | **6.17E-03** | 0.23 | 35.72 | 45.76 | -14.86 |
| GloVe | 6.66E-02 | **5.88E-04** | 0.24 | 31.33 | 41.37 | -12.66 |
| FastText | 5.72E-02 | **6.08E-04** | 0.24 | 31.39 | 41.43 | -12.7 |
| LSA | **4.41E-04** | **6.58E-03** | 0.23 | 35.83 | 45.87 | -14.92 |
| HAL | **1.74E-02** | **1.90E-03** | 0.23 | 33.54 | 43.58 | -13.77 |
| TaxonomicA | **3.86E-02** | **9.21E-04** | 0.24 | 32.18 | 42.22 | -13.09 |
| TaxonomicB | 7.43E-02 | **1.51E-03** | 0.24 | 33.1 | 43.14 | -13.55 |
| SimilarityR | 4.51E-01 | **1.99E-04** | 0.25 | 29.27 | 39.31 | -11.63 |
| AssociationR | **1.31E-02** | **1.66E-03** | 0.24 | 33.29 | 43.33 | -13.64 |
| Control12 | **1.63E-06** | 4.58E-01 | 0.2 | 42.78 | 52.82 | -18.39 |
| SimilarityR controlled for: |  |  |  |  |  |  |
| Exp48 | 2.91E-01 | **9.19E-03** | 0.27 | 22.94 | 32.99 | -8.47 |
| Lancaster9 | **2.22E-04** | **3.47E-02** | 0.26 | 25.31 | 35.35 | -9.65 |
| word2vec | **8.24E-04** | 8.37E-01 | 0.25 | 29.8 | 39.84 | -11.9 |
| GloVe | **1.59E-02** | 3.20E-01 | 0.25 | 28.84 | 38.88 | -11.42 |
| FastText | **1.66E-02** | 4.40E-01 | 0.25 | 29.24 | 39.28 | -11.62 |
| LSA | **1.58E-05** | 5.24E-01 | 0.25 | 29.43 | 39.47 | -11.72 |
| HAL | **1.62E-03** | 4.20E-01 | 0.25 | 29.18 | 39.23 | -11.59 |
| TaxonomicA | **3.56E-03** | 1.66E-01 | 0.25 | 27.89 | 37.93 | -10.95 |
| TaxonomicB | **6.20E-03** | 2.93E-01 | 0.25 | 28.72 | 38.76 | -11.36 |
| WordNet | **1.99E-04** | 4.51E-01 | 0.25 | 29.27 | 39.31 | -11.63 |
| AssociationR | **1.85E-03** | 6.99E-01 | 0.25 | 29.69 | 39.73 | -11.85 |
| Control12 | **3.36E-10** | 5.30E-02 | 0.26 | 26.04 | 36.08 | -10.02 |
| AssociationR controlled for: |  |  |  |  |  |  |
| Exp48 | 4.32E-01 | **6.93E-05** | 0.27 | 23.45 | 33.49 | -8.73 |
| Lancaster9 | **1.48E-03** | **1.21E-03** | 0.25 | 28.9 | 38.94 | -11.45 |
| word2vec | **1.06E-02** | **2.65E-02** | 0.23 | 34.54 | 44.58 | -14.27 |
| GloVe | 1.21E-01 | **7.61E-03** | 0.24 | 32.3 | 42.35 | -13.15 |
| FastText | 8.96E-02 | **6.96E-03** | 0.24 | 32.14 | 42.18 | -13.07 |
| LSA | **4.93E-04** | 6.34E-02 | 0.22 | 36.04 | 46.08 | -15.02 |
| HAL | **1.90E-02** | **1.64E-02** | 0.23 | 33.69 | 43.73 | -13.84 |
| TaxonomicA | **1.16E-02** | **2.24E-03** | 0.25 | 30.05 | 40.09 | -12.02 |
| TaxonomicB | **1.75E-02** | **3.02E-03** | 0.24 | 30.6 | 40.64 | -12.3 |
| WordNet | **1.66E-03** | **1.31E-02** | 0.24 | 33.29 | 43.33 | -13.64 |
| SimilarityR | 6.99E-01 | **1.85E-03** | 0.25 | 29.69 | 39.73 | -11.85 |
| Control12 | **2.96E-07** | 7.41E-01 | 0.21 | 39.44 | 49.48 | -16.72 |
| Control12 controlled for: |  |  |  |  |  |  |
| Exp48 | **1.71E-04** | **9.30E-14** | 0.32 | 9.71 | 19.75 | -1.85 |
| Lancaster9 | 3.57E-01 | **1.67E-07** | 0.22 | 38.31 | 48.35 | -16.15 |
| word2vec | 9.38E-01 | **7.17E-07** | 0.21 | 41.17 | 51.21 | -17.59 |
| GloVe | 6.34E-01 | **2.44E-08** | 0.23 | 34.52 | 44.56 | -14.26 |
| FastText | 5.47E-01 | **2.68E-08** | 0.23 | 34.7 | 44.74 | -14.35 |
| LSA | 1.88E-01 | **1.18E-05** | 0.18 | 46.63 | 56.67 | -20.31 |
| HAL | 9.58E-01 | **2.74E-07** | 0.21 | 39.28 | 49.33 | -16.64 |
| TaxonomicA | 6.93E-01 | **6.24E-08** | 0.22 | 36.37 | 46.41 | -15.19 |
| TaxonomicB | 4.33E-01 | **4.47E-08** | 0.23 | 35.71 | 45.76 | -14.86 |
| WordNet | 4.58E-01 | **1.63E-06** | 0.2 | 42.78 | 52.82 | -18.39 |
| SimilarityR | 5.30E-02 | **3.36E-10** | 0.26 | 26.04 | 36.08 | -10.02 |
| AssociationR | 7.41E-01 | **2.96E-07** | 0.21 | 39.44 | 49.48 | -16.72 |

**Supplemental Table 10.**
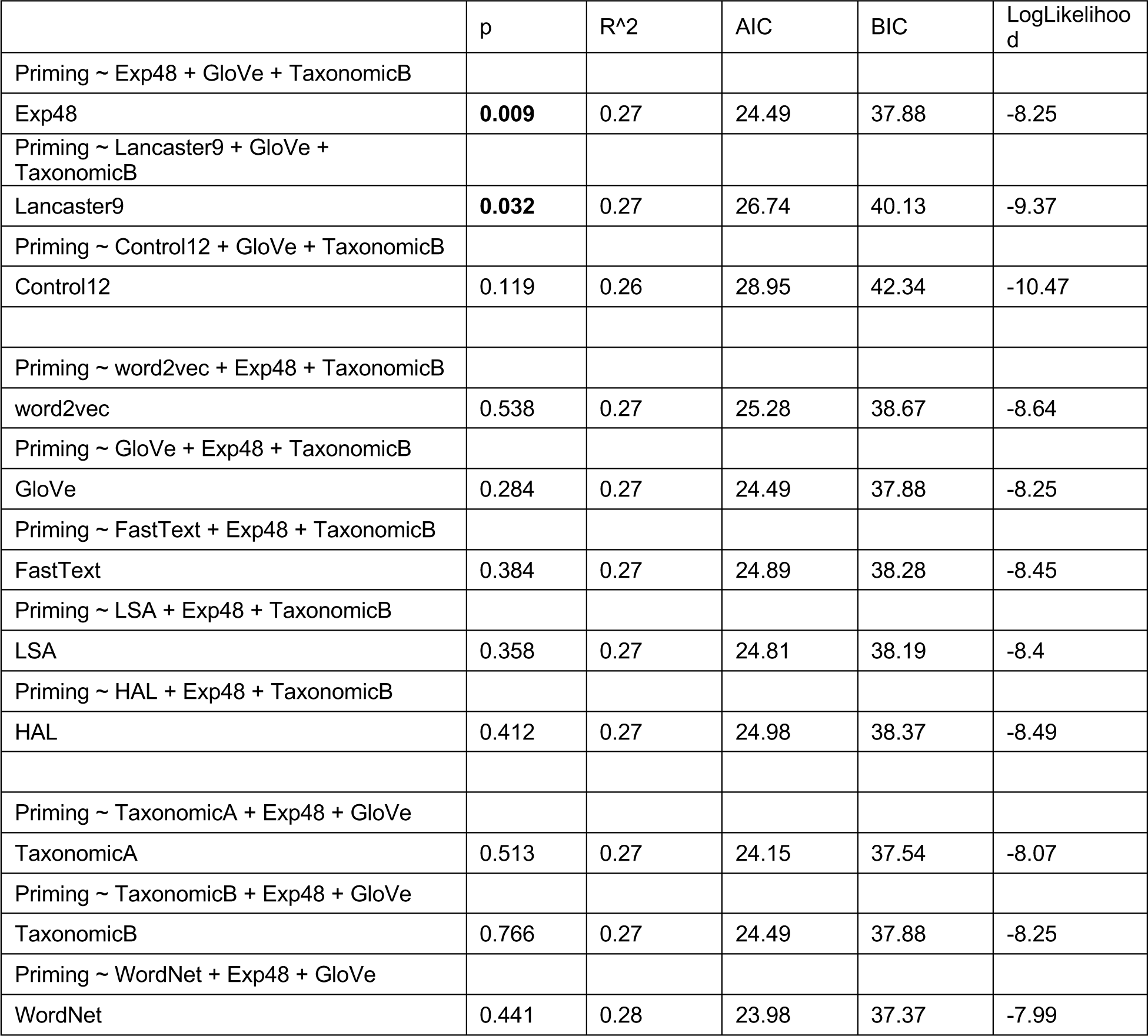
Results of regression models including three regressors at a time. Predictions from each semantic model were simultaneously controlled for the predictions of the top-performing semantic model of each of the other two types.

## Notes

### Competing Interest Statement

The authors have declared no competing interest.

### Summary of Updates

New analyses were added (4 new representational models and supplemental linear model analyses), the Introduction was revised to clarify the rationale for the analyses, and the Discussion was extended to address theoretical and methodological issues more thoroughly.

